# ProSiteHunter: A unified framework for sequence-based prediction of protein-nucleic acid and protein-protein binding sites

**DOI:** 10.1101/2025.10.22.683834

**Authors:** Dongliang Hou, Qihang Zhen, Zexin Lv, Xinyue Cui, Suhui Wang, Minghua Hou, Zhan Zhou, Xiaogen Zhou, Guijun Zhang

## Abstract

Accurate prediction of protein binding sites is essential for elucidating protein function, understanding molecular interaction mechanisms, and facilitating drug design. However, existing sequence-based approaches are often designed for specific binding-site types and therefore lack generality, whereas structure-based methods typically rely on high-quality structural models, limiting their applicability. Here, we introduce ProSiteHunter, a unified sequence-based framework for protein binding-site prediction, which integrates a fine-tuned protein language model (SiteT5) with a multi-source feature-fusion network that incorporates evolutionary, geometric, and statistical features, while employing bidirectional semantics, local associations, and global dependencies for comprehensive binding-site characterization. The method was systematically evaluated on diverse binding sites prediction tasks, where ProSiteHunter achieved a 39.1% average improvement in PRAUC for protein-DNA/RNA/protein tasks and a 7.4% PRAUC enhancement on the particularly challenging antibody-antigen task over state-of-the-art methods. Moreover, ProSiteHunter is capable of identifying local flexible sites that complement AlphaFold3 predictions and improving the accuracy of antibody-antigen interaction prediction. These results highlight ProSiteHunter as an efficient and unified approach for accurate and robust prediction of diverse protein binding sites.

## Introduction

The interactions between proteins and macromolecules are the core mechanisms of life activities, playing an important role in processes such as cell signal transduction, gene expression regulation, and immune response^1-3^. The binding sites represent critical regions that mediate interactions between these biomolecules. Accurate determination of these sites is essential not only for elucidating protein function but also for providing key insights into drug target identification, mechanistic studies of biomolecular processes, and antibody design^4-6^. The investigation of protein-nucleic acid binding sites contributes to elucidating gene regulatory networks^7,8^ and the mechanisms of transcriptional regulation^9^. Similarly, the prediction of protein-protein binding sites can help uncover signaling pathways associated with diseases^1^. The identification of antibody-antigen binding sites, as a distinct category of protein interactions, holds particular research significance. It provides a critical foundation for epitope-centric vaccine design^10^ and, importantly, for the development of antibody-based therapeutics^11^. Traditional biological experimental methods (such as affinity purification-mass spectrometry^12^, immunoprecipitation^13^ and hybrid screening^14,15^), while capable of precisely detecting interaction sites, are time-consuming and costly, and have stringent requirements for experimental conditions, making it difficult to meet the needs of large-scale research. Consequently, the development of efficient and accurate computational methods for predicting binding sites has emerged as a critical scientific challenge demanding urgent solutions.

Structure-based computational methods are mainstream approaches that are generally divided into two groups: graph neural network (GNN) based methods and geometric feature engineering-based methods. GNN-based methods model the spatial topological relationships between residues by constructing a graph representation of protein structures. Typical works include: GraphPPIS^16^, which predicts protein interaction sites using a graph network constructed from secondary structure and residue contact information; GraphBind^17^, which integrates physicochemical features and hierarchical graph networks to predict nucleic acid binding residues; and EGPDI^18^, which constructs a graph convolutional network based on residue coordinates to predict protein-DNA binding sites. Geometric feature engineering methods focus on extracting key features from three-dimensional spatial information. Representative methods include PeSTo^19^, which predicts protein interaction sites by characterizing protein surface shapes with atomic point clouds; epitope3D^20^, which describes residue neighbors based on the composition of residues at different distances to predict conformational B cell epitopes; and SEMA2.0-3D^21^, which uses 3Di structural encoding to achieve epitopes prediction. While these methods have achieved remarkable success in predictive performance, several important challenges remain to be addressed for their broader practical application. A primary limitation is the strong dependency on high-quality structural data, where even minor conformational deviations can lead to prediction inaccuracies. Although advances in computational tools such as AlphaFold2^22^ have dramatically improved protein backbone prediction, accurately modeling flexible binding regions remains a major technical hurdle^23^. Furthermore, many functional interactions rely on dynamic conformational changes^24^, suggesting that predictions derived from static structural snapshots may fail to capture authentic binding mechanisms^25^. These inherent constraints currently limit the generalizability and practical utility of structure-based binding site prediction methods.

Sequence-based prediction methods have emerged as a prominent and rapidly advancing frontier in bioinformatics, requiring only protein primary sequences as input. This approach demonstrates significantly greater robustness than structure-based methods, as it circumvents conformational prediction inaccuracies and inherent challenges posed by protein structural dynamics. Furthermore, sequence-based techniques substantially reduce computational resource requirements compared to structure-dependent approaches. Notably, recent advances in large language models have dramatically improved the accuracy of sequence-based binding site prediction^26,27^, further establishing this paradigm as a pivotal direction for future research. Sequence-based computational methods can be generally classified into three categories based on the predicted binding site types: the first category includes protein-protein interaction (PPI) sites prediction methods, where DELPHI^28^ predicts interactions by introducing protein binding barrier regions and PSSM matrices, and EnsemPPIS^29^ enhances prediction accuracy by integrating the ProtBERT large language model with a Transformer architecture; the second category involves protein-nucleic acid binding sites prediction methods, such as iDRNA-ITF^30^, which uses an inductive transfer framework to predict DNA/RNA binding residues, and CLAPE^31^, which employs contrastive learning to identify protein-nucleic acid binding sites; the third category comprises B-cell epitope prediction methods, including BepiPred-3.0^32^, which integrates the ESM-2^26^ language model for epitopes prediction, CALIBER^33^, which predicts linear and epitopes based on Bidirectional Long Short-Term Memory (BiLSTM) networks, and SEMA2.0-1D^21^, which optimizes epitopes feature learning for prediction by fine-tuning the ESM-2^26^ model.

While sequence-based binding site prediction methods have made remarkable progress, two critical challenges remain: limited task generalizability and inadequate modeling of sequence dependencies. Most existing approaches are tailored to specific types of interactions, such as protein-protein or protein-nucleic acid binding, using distinct feature sets and network architectures. Although this specialization enhances performance on dedicated tasks, it considerably restricts model generalization across different prediction scenarios. Furthermore, owing to constraints in local receptive fields and insufficient global context modeling, current methods often fail to capture long-range dependencies within sequences, a fundamental requirement for accurately identifying distal functional residues and achieving a comprehensive understanding of protein function mechanisms. Addressing these limitations would significantly advance the development of universal binding site predictors and provide a new paradigm for systematically deciphering the functional code of proteins.

In this work, we present ProSiteHunter, a universal sequence-based framework for predicting general binding sites between protein-nucleic acid and protein-protein interactions. The innovation of the proposed framework is manifested at three distinct levels. At the architectural design level, we have designed a multi-source feature fusion module that achieves spatial alignment and collaborative optimization of multimodal feature representation through three-track interaction. At the feature representation level, we integrated three complementary types of protein information: evolutionary features, statistical features, and geometric features to construct a multi-dimensional representation space of binding sites. At the sequence encoding level, we have developed SiteT5, a protein language model fine-tuned from ProtT5-XL-UniRef50^27^ by integrating evolutionary information. Compared to existing methods, ProSiteHunter not only exhibits broad applicability, capable of predicting protein-nucleic acid (DNA/RNA), protein-protein, and specialized protein (antibody-antigen) binding sites, but also demonstrates superior prediction accuracy and robustness. Cross-task validation further confirms that the integrated multi-source features and the unique architectural design are critical contributions to the model’s performance.

## Results

### Overview of ProSiteHunter

ProSiteHunter leverages a unified deep learning framework based on an encoder-decoder architecture. It takes a single amino acid sequence as input and outputs residue-level binding propensity scores, indicating the likelihood of each residue being part of a binding site. We employ an information enhancement strategy through multi-level feature optimization. Our framework predicts binding sites for four major types of protein interactions: protein-DNA, protein-RNA, protein-protein, and antibody-antigen. Four specialized models are independently trained on type-specific datasets, enabling effective identification of distinctive structural features and binding patterns pertinent to each interaction type.

ProSiteHunter comprises two main stages. In the feature fusion stage, a Multi-Source Feature Fusion (MSFF) with three-track semantic parsing is constructed to realize the complementary representation of sequence information (**Fig. 1a** illustrates the architectural optimization of the three networks, which share the same sequence representation (sr) as input.): (1) Multi-Scale CNN module, which uses serial convolutional kernels to extract short-range inter-residue information in the local receptive field; (2) BiLSTM is used for bidirectional semantic modeling, and directional sensitivity features are captured through forward and reverse sequence scanning; (3) The Gated Attention module learns global dependencies and explicitly captures long-distance residual associations. In particular, trainable weight matrices are employed to adaptively weight the features from the three tracks and project them into the Query (Q), Key (K), and Value (V) vectors for the cross-attention module, enabling dynamic alignment and fusion of the feature spaces. In the self-interaction stage, these representations are processed by a Multi-Level Interactive Learning (MIL) module composed of three blocks. Within this module, relevant information about binding sites is further refined through multi-head sub-attention with gating mechanisms. Nonlinear transformation and feature enhancement are then applied via a Position-wise Feed-Forward Network (PFFN). Finally, a classifier produces a probability score for each residue, indicating its likelihood of being a binding site.

**Fig. 1.**
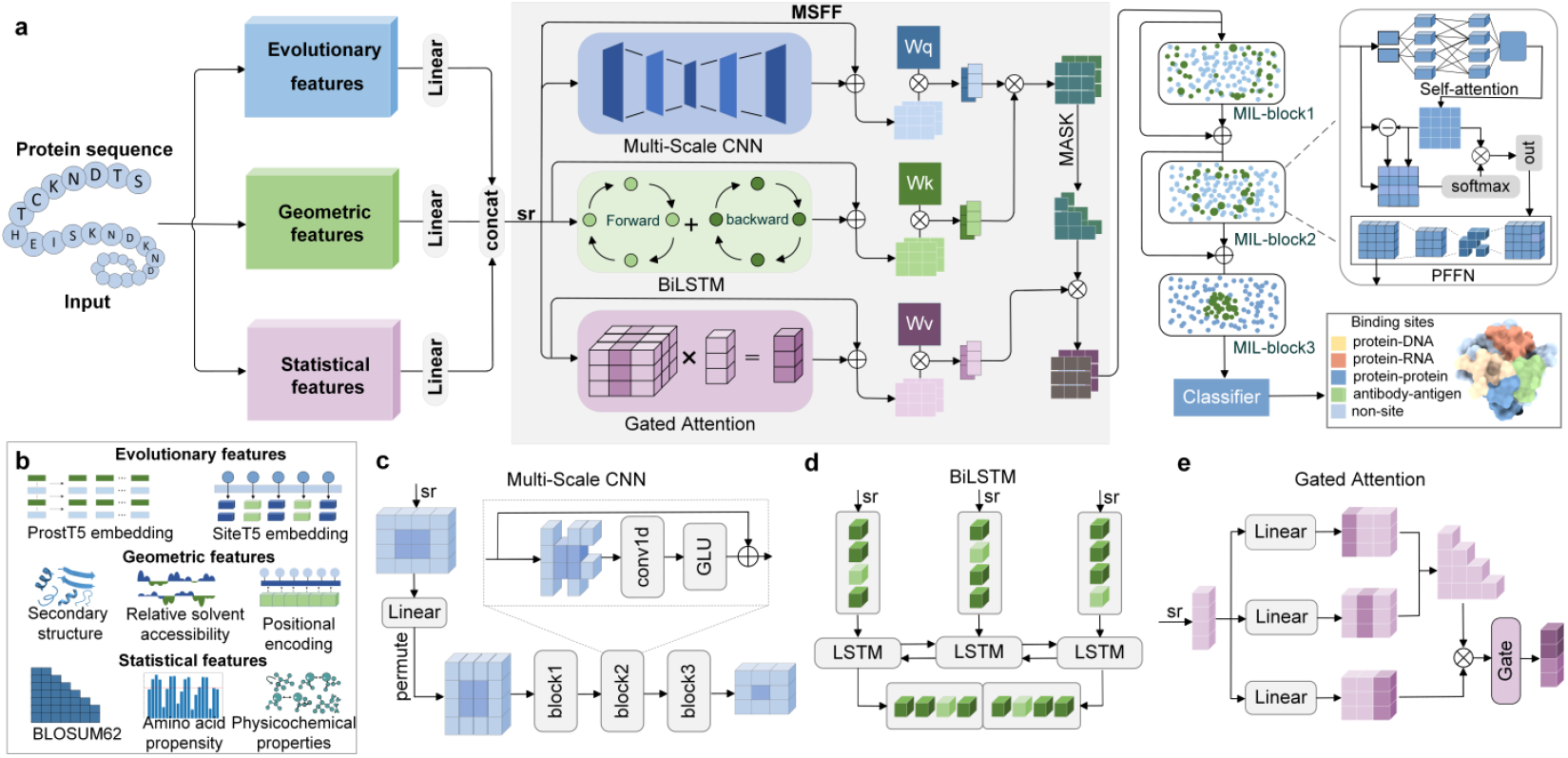
Pipeline of ProSiteHunter. **a**, ProSiteHunter takes a protein sequence as input and first extracts and integrates evolutionary, geometric, and statistical features to construct a multi-level sequence representation. The fused features are then passed into the Multi-Source Feature Fusion (MSFF) module, which consists of a Bidirectional Long Short-Term Memory (BiLSTM) module, a Multi-Scale Convolutional module (Multi-Scale CNN), and a Gated Attention module. The tensors generated by these three networks are mapped to Queries (Q), Keys (K), and Values (V), respectively, and integrated through a cross-attention mechanism to capture both local patterns and global dependencies. The fused representation is subsequently fed into the Multi-Layer Interaction Learning (MIL) module, which is composed of three stacked sub-blocks, each containing a multi-head self-attention unit, a gating unit, and a position-wise feed-forward network (PFFN), enabling progressive modeling of semantic interactions among residues. Finally, the model outputs the probability of each residue being part of a protein-DNA, protein-RNA, protein-protein, or antibody-antigen binding site through a classifier. The protein structure shown in the figure is a schematic diagram. **b**, Feature extraction integrates three complementary feature types: evolutionary, geometric, and statistical, to construct a multidimensional representation of each site. All features are collectively referred to as sequence representations (sr). **c**, The network of the Multi-Scale CNN module. **d**, The network of the BiLSTM module. **e**, The network of the Gated Attention module.

In the process of feature extraction, we leverage two Protein Large Language Models (PLLM), SiteT5 and ProstT5, to encode key features of proteins from the perspectives of evolutionary conservation and geometric plausibility, thereby generating significant synergistic effects in binding site prediction tasks. Among them, SiteT5 is a specialized model we developed by fine-tuning ProtT5-XL-UniRef50^27^, aiming to effectively capture the evolutionarily conserved features of functional sites. For four different binding site prediction tasks, we searched for multiple sequence alignments (MSAs) from the protein sequences of the corresponding training sets and selected sub-MSAs as inputs, enabling the model to learn the distinctive patterns and features of different protein families. Beyond the protein language model, we integrated three geometric features: symmetrically normalized position encoding, secondary structure, and relative solvent accessibility. These features quantify spatial exposure and structural context and provide a foundation for binding site localization. We further incorporated three statistical features: BLOSUM62^34^ matrix, physicochemical properties, and amino acid propensity to characterize residue-level attributes. See the **‘Sequence feature’ section in Methods** and **Supplementary Table 1**.

By organically integrating multi-source feature synergy with a dynamic fusion mechanism, ProSiteHunter achieves a significant enhancement in the accuracy of protein binding site identification, all while maintaining a lightweight model architecture. This capability is substantiated by its robust performance in two challenging scenarios: structural inaccuracies from AlphaFold3 predictions and conformational changes in proteins driven by dynamics (**Figs. 2e and 5e**).

**Fig. 2.**
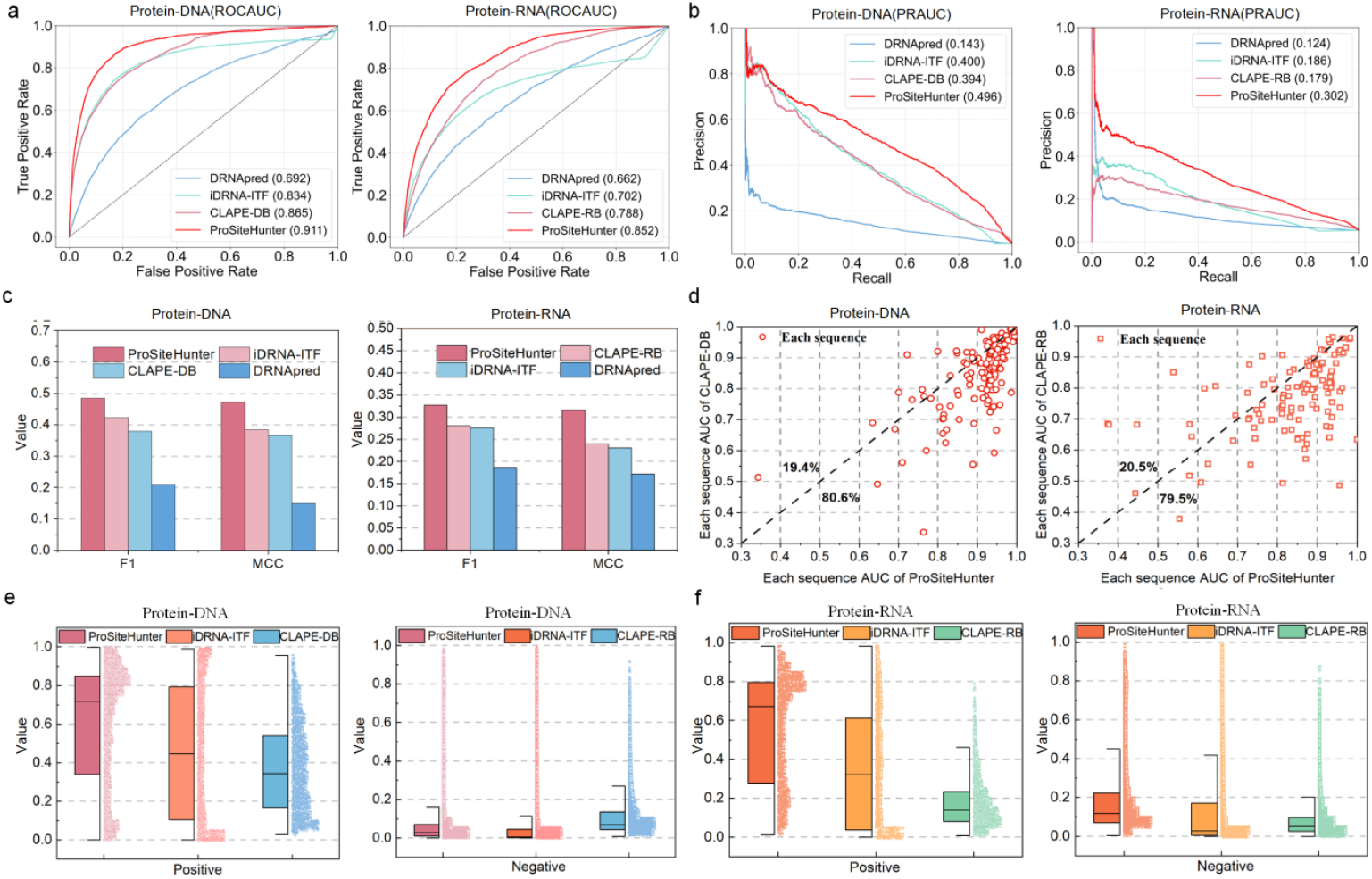
Performance of ProSiteHunter in protein-nucleic acid binding sites prediction. **a**, ROCAUC curve of ProSiteHunter on protein-nucleic acid prediction task. **b**, PRAUC curves of ProSiteHunter on protein-nucleic acid prediction tasks. **c**, The F1 and MCC performance comparison of the four methods: ProSiteHunter, CLAPE-DB, CLAPE-RB, iDRNA-ITF, and DRNApred. **d**, Scatter plots display the sequence-wise AUC values of ProSiteHunter versus CLAPE-DB for protein-DNA binding and CLAPE-RB for protein-RNA binding. **e**, Half-box plots comparing the classification performance of ProSiteHunter, CLAPE-DB, and iDRNA-ITF on positive and negative samples in protein-DNA binding site prediction. **f**, Half-box plots comparing the classification performance of ProSiteHunter, CLAPE-RB, and iDRNA-ITF on positive and negative samples in protein-RNA binding site prediction.

### Performance of protein-nucleic acid binding sites prediction

To fairly compare the predictive performance of ProSiteHunter on protein-nucleic acid binding sites, both we and the benchmark methods trained and tested our models using two different datasets from GraphBind^17^: one for protein-DNA and another for protein-RNA binding sites. In order to mitigate potential data leakage, a temporal split was applied by assigning all protein chains released before January 6, 2016 to the training set, and those released on or after that date to the test set. Furthermore, to address the class imbalance between positive and negative samples, protein chains were clustered. Within each cluster, binding site annotations were transferred to the longest chain to enhance the representativeness of the data.^51^ For further details, see the **‘Data set’ section in Methods**.

We benchmarked our method against state-of-the-art predictors, including DRNApred^35^, iDRNA-ITF^30^, CLAPE-DB^31^ and CLAPE-RB^31^. The predictions from DRNApred^35^ and iDRNA-ITF^30^ were generated using their respective web servers, while those for CLAPE-DB^31^ and CLAPE-RB^31^ were reproduced based on their publicly available source code^31^. The performance was evaluated using a comprehensive set of standard metrics, comprising the area under the receiver operating characteristic curve (ROCAUC), F1-score, and Matthew’s correlation coefficient (MCC), It is worth noting that in order to avoid the bias caused by the imbalance of positive and negative sample data in the test set (see **Supplementary Table 2**), we introduced the area under the precision-recall curve (PRAUC) to further evaluate ProSiteHunter performance.

As illustrated in **Fig. 2a**, ProSiteHunter demonstrates significantly superior performance compared to other benchmarked methods. Specifically, it achieved ROCAUC values of 0.911 and 0.852 for predicting protein-DNA and protein-RNA binding sites, respectively, surpassing the second-best method, CLAPE-DB^31^, by 5.3% and CLAPE-RB^31^ by 8.4%. Notably, ProSiteHunter also showed substantial advantages in other core metrics such as PRAUC, F1-score, and MCC (**Figs. 2b, 2c**). For protein-DNA binding site prediction, it improved by 25.9% in PRAUC, 21.6% in F1-score, and 28.9% in MCC compared to CLAPE-DB^31^. Similarly, for protein-RNA prediction, the corresponding improvements over CLAPE-RB reached 66.7% in PRAUC, 16.4% in F1-score, and 31.7% in MCC (**Supplementary Tables 4 and 5**). These results collectively indicate that ProSiteHunter maintains exceptional predictive accuracy even under highly imbalanced classification conditions, where the site-to-non-site ratio approaches 1:10 (**Supplementary Table 2**), highlighting its robust classification capability.

In the sequence-wise comparative analysis, ProSiteHunter consistently outperformed CLAPE-DB^31^ and CLAPE-RB^31^, achieving higher AUC values on 80.6% and 79.5% of test sequences in protein-DNA and protein-RNA binding site prediction, respectively (**Fig. 2d**). A similar trend was observed in comparisons with iDRNA-ITF^30^ and DRNApred^35^, where ProSiteHunter also demonstrated clear advantages (**Supplementary Fig. 1**). We hypothesize that this robust performance stems from our three-track feature fusion architecture, which synergistically integrates Multi-Scale CNN, BiLSTM, and a Gated Attention module. This architectural design, as further validated by the ablation studies in **Fig. 4c**, effectively combines site and non-site information across diverse representation spaces, thereby enhancing both discriminative power and generalization capacity. In contrast, approaches relying on single-network architectures may not sufficiently capture the complex and hierarchical determinants of binding sites, which could partially explain their comparatively limited predictive performance.

To evaluate the discriminatory ability of ProSiteHunter between binding sites and non-binding sites, we analyzed the predictive score distributions for both positive and negative samples across protein-DNA and protein-RNA tasks **(Figs. 2e, 2f)**. ProSiteHunter demonstrated a pronounced advantage over state-of-the-art methods such as iDRNA-ITF^30^, CLAPE-DB^31^, and CLAPE-RB^31^. Specifically, it assigned high prediction scores (mostly above 0.75) to binding sites, indicating strong discriminative confidence for positive samples. In comparison, baseline methods yielded predominantly lower scores (below 0.5) for true binding residues, reflecting limited sensitivity in identifying positive instances. These results confirm the superior capability of ProSiteHunter in recognizing binding sites and its robustness in class-imbalanced prediction scenarios.

ProSiteHunter demonstrates robust and consistent superiority over existing baseline methods across three fundamental aspects: comprehensive evaluation metrics, sequence-wise predictive accuracy, and discriminative capability in distinguishing binding from non-binding residues. We attribute this advantage to two core designs: the SiteT5 module, a protein language model-based encoder that captures rich evolutionary and biochemical information, and a three-track fusion architecture that integrates multi-scale and hierarchical sequence semantics. Ablation experiments (**Figs. 4c, 4f**) further confirm that both components contribute significantly to the model’s enhanced performance. The SiteT5 encoder provides highly discriminative residue-level features, while the fusion mechanism enables effective integration of multi-source contextual information. Collectively, these designs endow ProSiteHunter with strengthened generalization capacity and improved robustness in capturing the complex sequence features of protein-nucleic acid binding sites.

### Performance of protein binding sites and antigen epitopes prediction

To evaluate the applicability of the ProSiteHunter framework for predicting protein-protein interaction sites, we maintained its original architecture and performed training and validation using the EnsemPPIS^29^ dataset, which contains 422 protein sequences with less than 25% sequence identity. Further details are described in the **‘Data set’ section of Methods**. Benchmark results showed that ProSiteHunter maintained strong predictive performance on this task. Given the importance of antibody-antigen interactions in pharmaceutical applications, we further assessed the performance of ProSiteHunter on antigen epitopes prediction. To ensure fair comparison, we trained and tested the model on the SEMA1.0^36^ antibody-antigen conformational epitope dataset (as epitope labels are masked in SEMA2.0^21^). This dataset assigns protein chains published before January 1, 2020, to the training set and protein chains published on or after that date to the test set. Further details are described in the **‘Data set’ section of Methods**. Benchmark results demonstrated that ProSiteHunter consistently outperformed other baseline methods.

In the protein binding sites prediction task (**Figs. 3a and 3b**), ProSiteHunter demonstrated significantly superior performance over baseline methods, including EnsemPPIS^29^, DELPHI^28^, and DeepPPISP^37^. Relative to the best-performing baseline, EnsemPPIS^29^, ProSiteHunter achieved notable improvements across multiple evaluation metrics, with increases of 9.3% in ROCAUC, 24.7% in PRAUC, 16.1% in F1-score, and 34.3% in MCC (**Supplementary Table 6**). Similarly, in the antigen epitope prediction task, ProSiteHunter consistently outperformed the top-performing baseline, CALIBER^33^ (**Figs. 3a and 3b**), achieving improvements of 2.1%, 7.4%, 8.3%, and 10.0% in ROCAUC, PRAUC, F1-score, and MCC, respectively (**Supplementary Table 7**). Collectively, these results highlight that ProSiteHunter delivers high predictive accuracy while maintaining strong generalization capability across diverse prediction tasks, underscoring its effectiveness as a unified framework for binding site prediction.

**Fig. 3.**
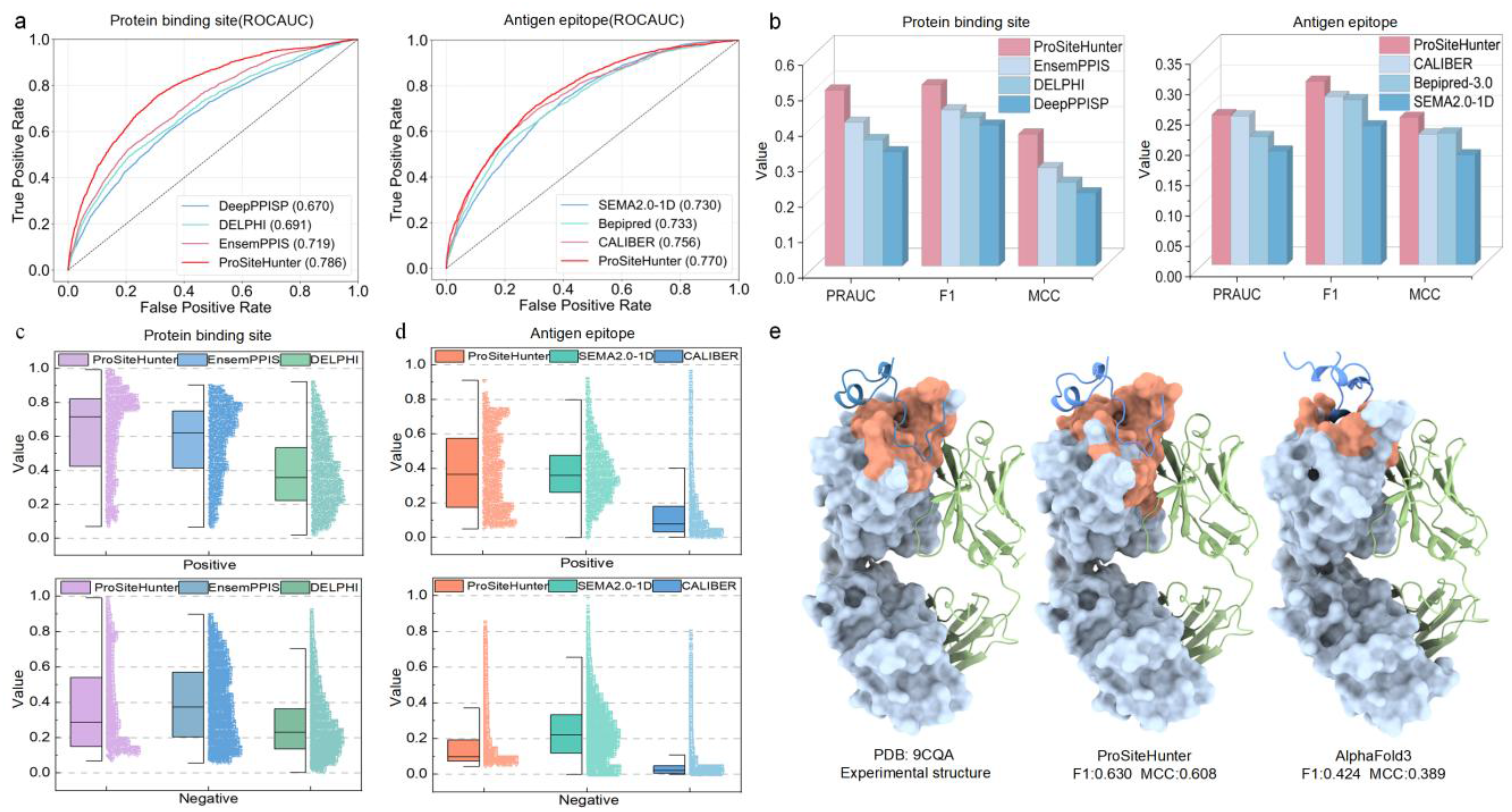
Performance of ProSiteHunter in protein binding sites, antigen epitopes prediction. **a**, ROCAUC curves of ProSiteHunter for protein binding site and antigen epitope prediction tasks. **b**, The PRAUC, F1 and MCC performance comparison of the four methods: ProSiteHunter, EnsemPPIS, DELPHI, and DeepPPISP. **c**, Half-box plots display the classification performance of ProSiteHunter versus EnsemPPIS and DELPHI on positive and negative samples in the protein binding sites prediction task. **d**, Half-box plots display the classification performance of ProSiteHunter versus SEMA2.0-1D and CALIBER on positive and negative samples in the antigen epitope prediction task. **e**, Comparison of binding site predictions for the CASP16 target T1225 (PDB ID: 9CQA): the experimental structure (left, the light blue portion represents the antibody heavy chain region, which binds to the dark blue respiratory syncytial virus glycoprotein protein. The orange region indicates the true binding sites), predictions from ProSiteHunter (middle; orange region for predicted binding sites), and predictions from AlphaFold3 (right; orange region for predicted binding sites).

In addition, we tested the classification performance of ProSiteHunter against baseline methods (EnsemPPIS^29^ and DELPHI^28^) (**Fig. 3c**). ProSiteHunter outperformed EnsemPPIS^29^, achieving 20.6% higher accuracy in recognizing positive samples. Furthermore, ProSiteHunter exhibited superior predictive confidence, with the majority of its positive sample scores concentrated in the 0.75-1.0 range, whereas DELPHI’s predictions were predominantly distributed across the lower 0.1-0.6 interval. These results demonstrate its strong discriminative power in identifying positive samples. In more challenging antigen epitopes prediction tasks, ProSiteHunter maintained robust classification performance (**Fig. 3d**). In contrast, other methods are overly conservative in predicting sites and are not sensitive to the distinction between positive and negative samples. These performance differences likely arise from the reliance of these methods on single-modal sequence features, without incorporating complementary information such as protein spatial topology, long-range interactions, or evolutionary conservation. Consequently, they face challenges in establishing precise decision boundaries when classifying positive and negative samples.

**Fig. 3e** presents the structural model of the complex between the central conserved domain of respiratory syncytial virus glycoprotein (RSV G) and monoclonal antibody 1G1, serving as a representative case study that demonstrates the capability of our approach in resolving functional epitope-paratope interfaces. RSV is a major pathogen causing severe lower respiratory tract infections in infants, young children, and the elderly. Its attachment glycoprotein facilitates viral entry into host cells and modulates the host immune response by binding to the chemokine receptor CX3CR1^38^. In the native protein structure shown on the left side of **Fig. 3e**, the light blue portion represents the antibody heavy chain region, which binds to the dark blue RSV G protein. The orange region indicates the actual binding sites. This protein, designated as T1225 in CASP16. We used ProSiteHunter to predict binding sites on antibody heavy chains, achieving an F1 score of 0.630 and an MCC of 0.608. Compared with AlphaFold3, which yielded an F1 score of 0.424 and an MCC of 0.389, this represents improvements of 48.6% and 56.3%, respectively. Structure-based methods often predict incorrect binding site regions when the predicted protein structure deviates from experimental results. In contrast, ProSiteHunter delivers superior and more reliable performance in identifying the majority of binding sites, establishing it as a highly effective complement to AlphaFold3^39^.

Our analysis confirms the consistent superiority of ProSiteHunter over benchmark methods across all four sub-tasks (protein-DNA, protein-RNA, protein-protein, and antibody-antigen), despite significant variations in intrinsic prediction difficulty. Notably, the task of antigen epitopes prediction is more challenging than protein-DNA binding sites prediction. We propose that the observed discrepancy arises from distinct binding mechanisms: antibodies primarily interact with epitopes via their complementarity-determining regions (CDRs), whose conformational diversity contributes to increased structural heterogeneity within epitopes. This observation aligns with the documented difficulties of both AlphaFold2^22^ and AlphaFold3^39^ in modeling antibody-antigen complexes. Moreover, these findings suggest that a hybrid strategy integrating sequence-based epitope prediction with the structural modeling capabilities of AlphaFold2/3 may be a worthwhile avenue to explore for improving the accuracy of antibody-antigen complex determination.

### Ablation studies and interpretability analysis

To validate the rationale of feature selection and the effectiveness of the network architecture in ProSiteHunter, we first performed feature correlation analyses and ablation studies (including both feature and network ablations). To further examine the model’s ability to identify binding sites and capture underlying interaction patterns, we conducted amino acid propensity analyses within binding regions and integrated them with t-SNE40 visualization. These analyses demonstrated that the selected features make significant contributions to binding site prediction, and that the designed architecture excels in feature integration and pattern learning, enabling effective discrimination between binding and non-binding residues. Collectively, these results enhance the interpretability of the model and substantiate its effectiveness in both binding site identification and pattern recognition.

To assess the interdependence among different features, we conducted a systematic correlation analysis using cosine similarity as the evaluation metric. In particular, we focused on SiteT5 and relative solvent accessibility (RSA), as SiteT5 represents our evolution-based language model, whereas RSA is a classical structural descriptor closely associated with binding site formation (**Figs. 4a, 4b**). ProstT5^41^ and SiteT5 showed a moderate correlation (45%), reflecting that although they share similarities, they have different focuses due to different fine-tuning strategies: ProstT5^41^ relies on Foldseek^42^ 3Di sequences for structure guidance, while SiteT5 focuses on evolutionary information. SiteT5 generally showed low correlations with traditional physicochemical features (isoelectric point, polarizability, etc.) (<20%), demonstrating the importance of a multi-feature fusion strategy. RSA showed weak correlations with SiteT5 (9%) and ProstT5^41^ (16%), highlighting the limitations of language models in capturing structural features and the necessity of incorporating RSA into sequence prediction. Isoelectric point (77%), polarizability (65%), and amino acid propensity (82%) were highly correlated with RSA, further highlighting the close connection between solvent accessibility and functional residues. Overall, correlation analyses reveal the complementary roles of each indispensable feature category: language models provide semantic and evolutionary constraints, physicochemical descriptors reveal residue-level properties, and geometric descriptors capture the spatial environment. Their synergistic integration is therefore crucial for accurate binding site prediction.

**Fig. 4.**
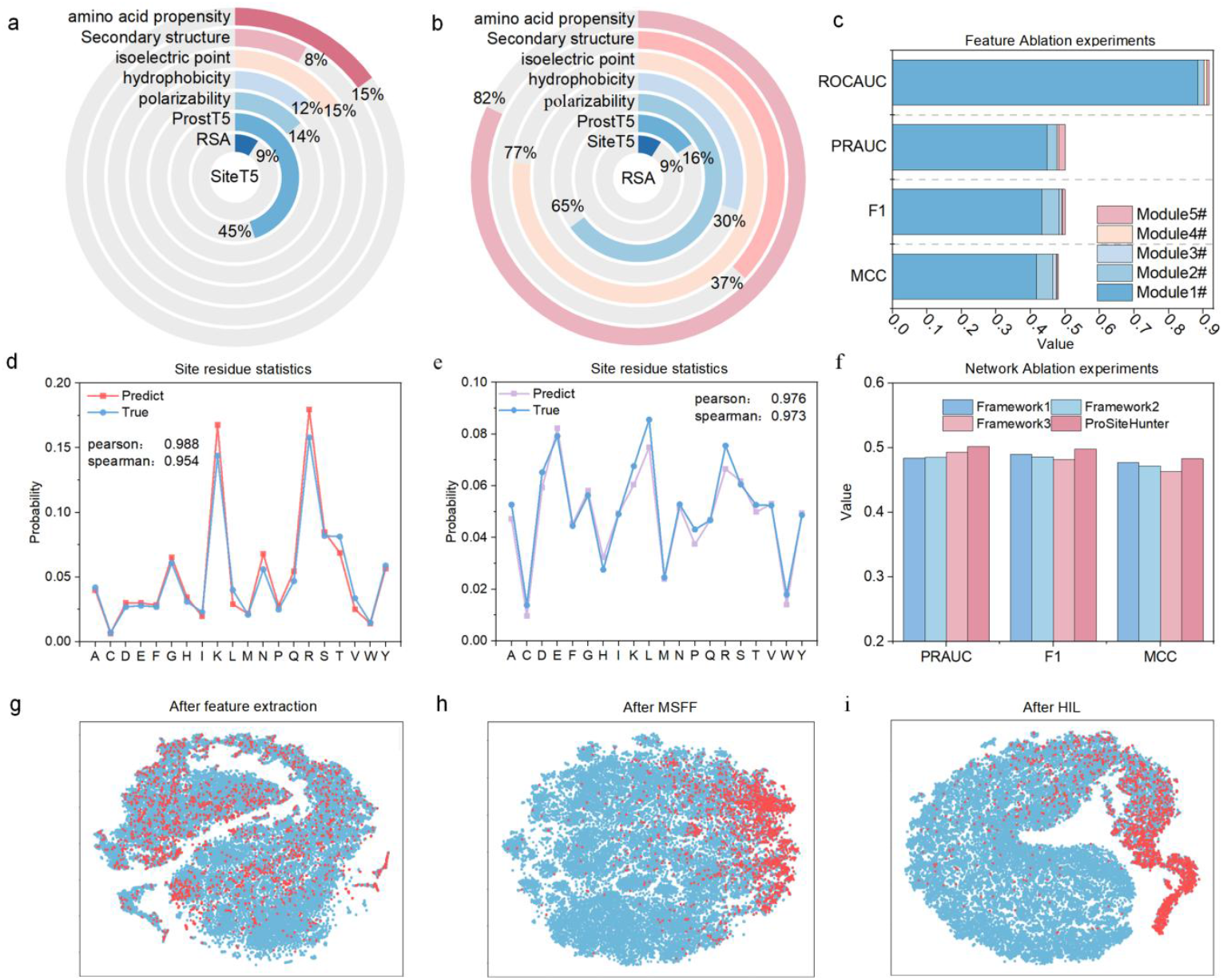
Ablation studies and interpretability analysis. **a**, Correlation analysis between SiteT5 embedding and a set of representative features, including RSA, ProstT5, polarizability, hydrophobicity, isoelectric point, secondary structure, and amino acid propensity. **b**, Correlation analysis between RSA embedding and a set of representative features, including SiteT5, ProstT5, polarizability, hydrophobicity, isoelectric point, secondary structure, and amino acid propensity. **c**, Characteristic ablation experiments. **d**, Amino acid propensity distribution of real protein-DNA binding sites and ProSiteHunter predicted binding sites. **e**, Amino acid propensity distribution of real protein-protein binding sites and ProSiteHunter predicted binding sites. **f**, Network ablation experiments. **g-i**, The t-SNE graph is used to map the high-dimensional tensor after feature extraction, MSFF module, and HIL module into a two-dimensional space to describe the relationship between protein-DNA binding sites (red) and non-sites (blue).

We then conducted systematic feature ablation experiments (**Fig. 4c**) to evaluate the contributions of different feature categories. Compared to a baseline module using only ProstT5^41^ (Module 1#), incorporating SiteT5 (Module 2#) improved MCC and F1 scores by over 11%, while improving PRAUC and ROCAUC by 6.3% and 2.2%, respectively. Geometric features primarily improved PRAUC, contributing an additional 4.2% gain from Module 4# to Module 5#. Detailed results are provided in **Supplementary Tables 1, 3**. Overall, these experiments demonstrated that the SiteT5 language model and geometric descriptors (positional encoding, secondary structure, and relative solvent accessibility) produced the most significant performance gains, confirming their key roles in accurate binding site prediction and providing empirical support for our multi-feature fusion strategy.

In order to systematically verify the superiority of the ProSiteHunter framework, we designed three comparison frameworks: Framework1 uses a BiLSTM encoder for self-attention decoders, Framework2 uses a Multi-Scale CNN encoder for self-attention decoders, and Framework3 builds an encoder-decoder structure based entirely on the self-attention mechanism. Experimental results show (**Fig. 4f**) that these three single semantic architectures have their own strengths and perform well in some indicators, but ProSiteHunter achieves the best in the three key indicators of PRAUC, F1 and MCC by integrating multi-semantic patterns, with an average improvement of 2.4%. This result not only proves the necessity of multi-source feature fusion, but also shows that the three-track semantic fusion strategy can effectively integrate the advantages of different architectures, thereby improving the overall performance of protein binding sites prediction.

To evaluate model interpretability (**Fig. 4g-4i**), we constructed a test set comprising 129 DNA-binding protein sequences, totaling 37,515 residues, including 2,240 binding sites (positive samples) and 35,275 non-binding sites (negative samples). In the visualizations, red points denote binding sites and blue points denote non-binding sites. At the feature extraction stage, binding sites were randomly scattered among non-binding sites. With the incorporation of the multi-source feature fusion module, binding sites progressively clustered toward the right region, and this trend became more pronounced after further processing by the interactive learning module. These results demonstrate that ProSiteHunter effectively distinguishes positive from negative samples, underscoring its advantage in binding site discrimination. The respective interpretability experiments for protein-RNA, protein-protein, and antibody-antigen binding sites are presented in **Supplementary Fig. 3**.

As a unified framework, ProSiteHunter demonstrates versatile capability by achieving robust performance in both protein-DNA and protein-protein binding mode prediction. As shown in **Fig. 4d**, the predicted amino acid frequency distribution of protein-DNA binding sites (red line) closely matches the ground truth (blue line) (Pearson = 0.988; Spearman = 0.954). Similarly, in protein-protein interaction prediction (**Fig. 4e**), the predicted distribution (purple line) aligns well with the experimental data (blue line) (Pearson = 0.976; Spearman = 0.973). These findings indicate that ProSiteHunter accurately captures specific binding patterns, owing to the deep representation power of its multi-source feature fusion module and the superior generalization enabled by its hierarchical feature extraction and dynamic fusion mechanisms. Collectively, our work suggests that ProSiteHunter could become a widely applicable tool for the study of diverse protein-macromolecule interactions. Further analysis (**Supplementary Fig. 2**) revealed two notable insights: first, protein-DNA and protein-RNA interactions share similar binding patterns, while protein-protein and antibody-antigen interactions also exhibit overlapping binding features; second, a higher proportion of loops in the binding-site region generally confers greater structural flexibility, thereby increasing the difficulty of accurate binding site prediction.

### Case study

To systematically evaluate the predictive performance of ProSiteHunter in different biological macromolecule interactions, we selected four representative proteins from the test set: PDB ID: 5H58 (protein-DNA binding, **Fig. 5a**), PDB ID:6D12 (protein-RNA binding, **Fig. 5b**), PDB ID: 3E1Z (protein-protein binding, **Fig. 5c**), and PDB ID: 7MJS (antibody-antigen binding, **Fig. 5d**). These cases exemplify key binding site characteristics: the interacting residues form tight structural clusters despite being discretely distributed in the linear sequence. The conformational dynamics of hen egg-white lysozyme (PDB: 4GN4/1J1X) are of particular interest, as conformational changes in its loop regions enable the formation of two distinct protein complexes (4GN4 and 1J1X) (**Fig. 5e**). The experimental results show that ProSiteHunter is able to effectively identify this change in binding sites region caused by protein conformational dynamics. The experimental results demonstrate ProSiteHunter’s capability for accurate identification of binding site alterations induced by protein conformational dynamics. This capability not only confirms its potential as a general-purpose prediction framework across diverse protein classes but also highlights its applicability in profiling dynamic binding sites.

**Fig. 5.**
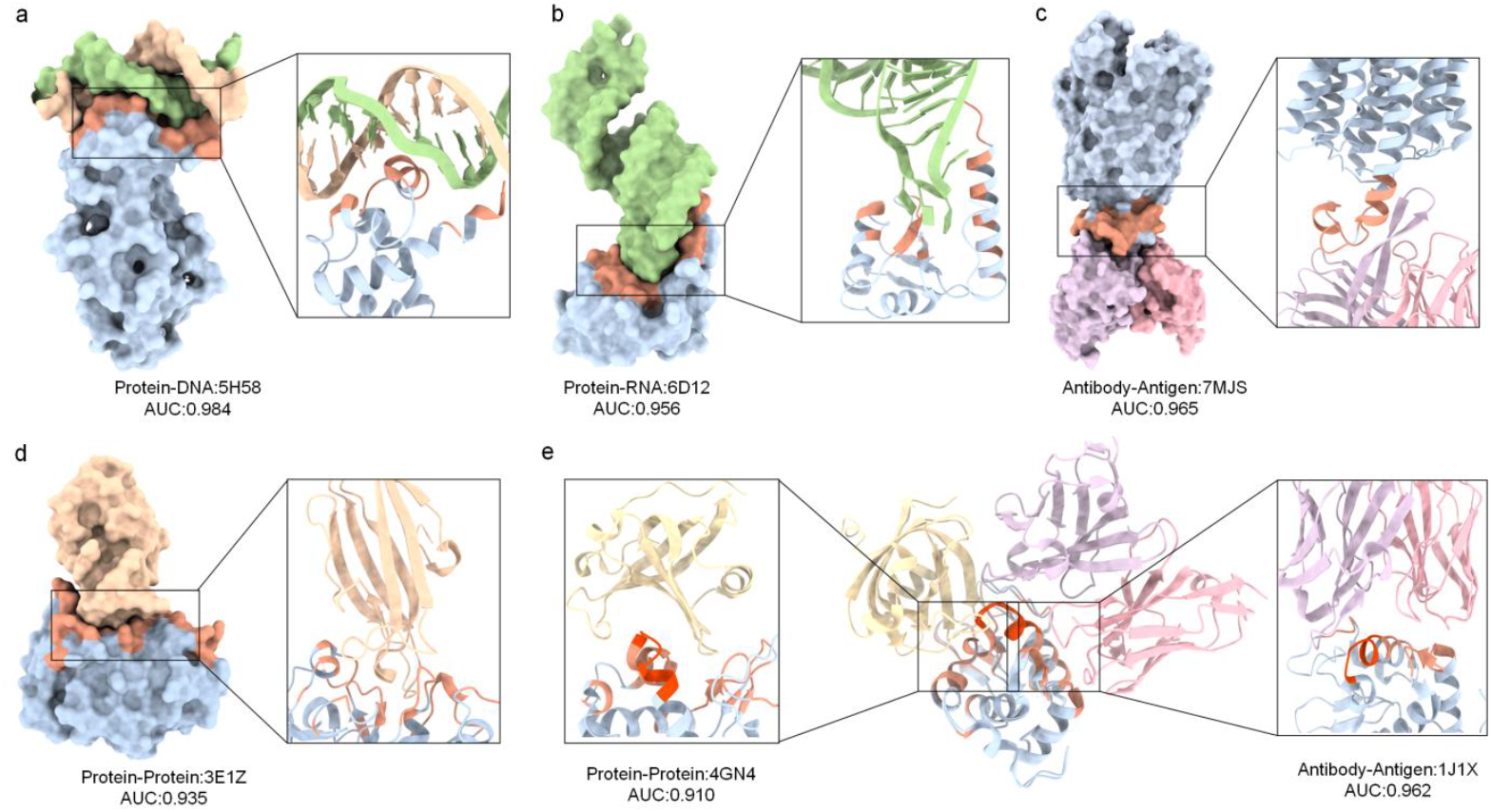
Special case of five different binding sites. **a**, The Protein (PDB ID:5H58) represents CprB (blue) from streptomyces coelicolor in complex with its cognate operator sequences (green and yellow). The orange region is the predicted binding sites. **b**, The Protein (PDB ID:6D12) represents the crystal structure of the human Larp7 (blue) C-terminal xRRM domain bound to the 7SK stem-loop 4 RNA (green). The orange region indicates the predicted binding sites. **c**, The Protein (PDB ID:7MJS) represents the complex of major promoter superfamily domain protein 2A (blue) and LPC-18:3 (pink and purple). The orange region indicates the predicted binding sites. **d**, The Protein (PDB ID:3E1Z) represents the crystal structure of the parasite protease inhibitor chagasin (yellow) in complex with papain (blue). The orange region indicates the predicted binding sites. **e**, The Protein (PDB ID:4GN4) represents AM2EP06 (yellow) bound to hen egg white lysozyme (blue). The Protein (PDB ID:1J1X) represents the crystal structure of the HyHEL-10 Fv mutant LS93A (purple, pink) bound to hen egg white lysozyme (blue). The predicted binding sites are highlighted in orange and red, and the red region indicates the overlap between the two binding sites.

5H58 is a complex of CprB of Streptomyces coelicolor and its biological longitudinal sequence^43^, in which tetracycline repressor family of transcription regulators (TetR-FTRs) transcriptional regulators regulate a variety of biological processes, including antibiotic biosynthesis, cell morphogenesis, and innate resistance. Among them, CprB is a member of TetR-FTRs from Streptomyces coelicolor, can specifically bind to operons (operons belong to DNA sequences, which are specific functional regions encoded by DNA in the genome of prokaryotes), and participates in the fine regulation of these physiological processes by regulating downstream gene expression^43^. ProSiteHunter accurately predicted most of the DNA binding sites on the protein (out of a total of 15 binding sites, 13 of which were successfully predicted. The prediction success rate was as high as 86.7%), and the AUC score reached 0.984, indicating that it has extremely high discriminant performance. As shown in **Fig. 5a**, the green and yellow models are the DNA double helix structure, the blue protein part is the predicted non-binding site, and the orange part is the predicted binding site region.

6D12 is a complex of the C-terminal xRRM domain of human Larp7 bound to the 7SK stem-loop 4 RNA^44^. The LARP superfamily is a diverse class of RNA-binding proteins involved in RNA processing, folding, and function. Larp7 binds to abundant long non-coding 7SK RNAs and is required for the assembly and function of 7SK ribonucleoprotein (RNP). Larp7 contains a C-terminal atypical RNA recognition motif (xRRM) that specifically binds to 7SK and P-TEFb assembly, and xRRM deletion may drive gastric cancer progression through epigenetic dysregulation such as aberrant transcriptional elongation^44^. As shown in **Fig. 5b**, the green part is the RNA model (forming a double helix structure, similar to DNA, but composed of adenine, guanine, cytosine, and uracil, and is a type of RNA), the blue part is the non-binding site area predicted using ProSiteHunter, and the orange area is the predicted binding site area, which tightly wraps RNA, with an AUC score of up to 0.956.

As shown in **Fig. 5c**, in the antibody-antigen complex 7MJS (PDB ID)^45^, partial loops and α-helical regions of the antigen (blue) form contacts with the antibody heavy chain (purple). Remarkably, ProSiteHunter accurately predicted these binding interface sites (orange), achieving high prediction performance in this case (AUC = 0.965). Antibody-antigen binding interfaces are often highly structurally diverse and flexible, making prediction challenging. However, ProSiteHunter reliably identified the antigen epitope in this case, further validating its versatility and effectiveness.

**Fig. 5d** (PDB ID: 3E1Z) shows the dimeric crystal structure of the parasitic protease inhibitor chagasin in complex with papain. The blue papain subunit contains multiple highly flexible loops that mediate binding to chagasin^46^. Although the conformational flexibility of these loops complicates binding site prediction and leads to some mislocalized predictions, ProSiteHunter successfully identified the majority of true binding sites, achieving an AUC-ROC value of 0.935 and demonstrating robust discriminatory performance.

As shown in **Fig. 5e**, we present a comparative analysis of two distinct complex structures of hen egg-white lysozyme (HEWL): the OBody AM2EP06-HEWL complex (PDB ID:4GN4^47^) and the HyHEL-10 Fv mutant LS93A-HEWL complex (PDB ID:1J1X^48^). These two different complexes are formed by a conformational change in their loop region, which results in a corresponding change in the binding sites region. ProSiteHunter accurately predicted the binding sites of egg white lysozyme in both complexes: the predicted AUC reached 0.910 in 4GN4 and 0.962 in 1J1X. This result demonstrates the potential of ProSiteHunter in predicting binding sites associated with conformational changes, and suggests that the dynamics of protein structure may be closely related to changes in binding sites regions. HEWL interacts specifically with different types of molecules, such as proteins or antibodies, through a conformational change in its loop region, which alters the region of the binding sites. This plasticity of binding sites, driven by conformational dynamics, may be one of the mechanisms by which proteins achieve functional diversification.

These results establish ProSiteHunter’s capability to accurately predict binding sites from sequence information alone. Notably, even its spurious predictions are informative, as they predominantly cluster spatially near genuine binding sites. This pattern strongly suggests that the model is implicitly learning the structural topology and evolutionary and physical constraints governing residue proximity directly from one-dimensional sequences (As illustrated in **Supplementary Fig. 4**). More remarkably, ProSiteHunter successfully predicts binding site remodeling induced by conformational changes from a single sequence (**Fig. 5e**), highlighting its potential to illuminate allosteric mechanisms and dynamic protein functions.

### Downstream application of ProSiteHunter: enhanced prediction of antibody-antigen interactions

Antibody-antigen interactions are a class of highly specific protein recognition processes that play a critical role in immune responses and biomedical applications^49^. The molecular basis of this interaction primarily relies on the precise recognition between the epitopes and the antibody complementarity-determining regions. To validate the applicability of ProSiteHunter in the field of immune recognition, this study specifically designed targeted downstream experiments: the epitopes information predicted by ProSiteHunter was integrated into MultisAAI^50^, an antibody-antigen interaction prediction framework developed in-house in our laboratory. The dataset constructed by MultisAAI^50^ includes 7580 antibody antigen pairs with a 1:1 ratio of positive to negative samples. We randomly selected 80% of the positive and negative samples as the training set and 20% as the test set. For the specific construction process of the dataset, please refer to MultisAAI^50^. Depending on whether to add sites features or not, we trained two models with the following results.

As shown in **Figs. 6a and 6b**, evaluation results based on ROCAUC, F1 score, precision, and recall indicate that incorporating epitope prediction features improves model performance by 5.74%, 7.20%, 7.48%, and 6.94%, respectively. Further comparative analysis (**Figs. 6c and 6d**) shows that this enhancement not only substantially increases the identification rate of positive samples (by 14.1%, **Fig. 6c**) but also strengthens the discrimination of negative samples (by 5.2%, **Fig. 6d**). Together, these findings underscore the critical role of epitope information in predicting antibody-antigen interactions. ProSiteHunter effectively pinpoints most epitopes, and the site information it provides further enhances both the reliability and interpretability of interaction predictions.

**Fig. 6.**
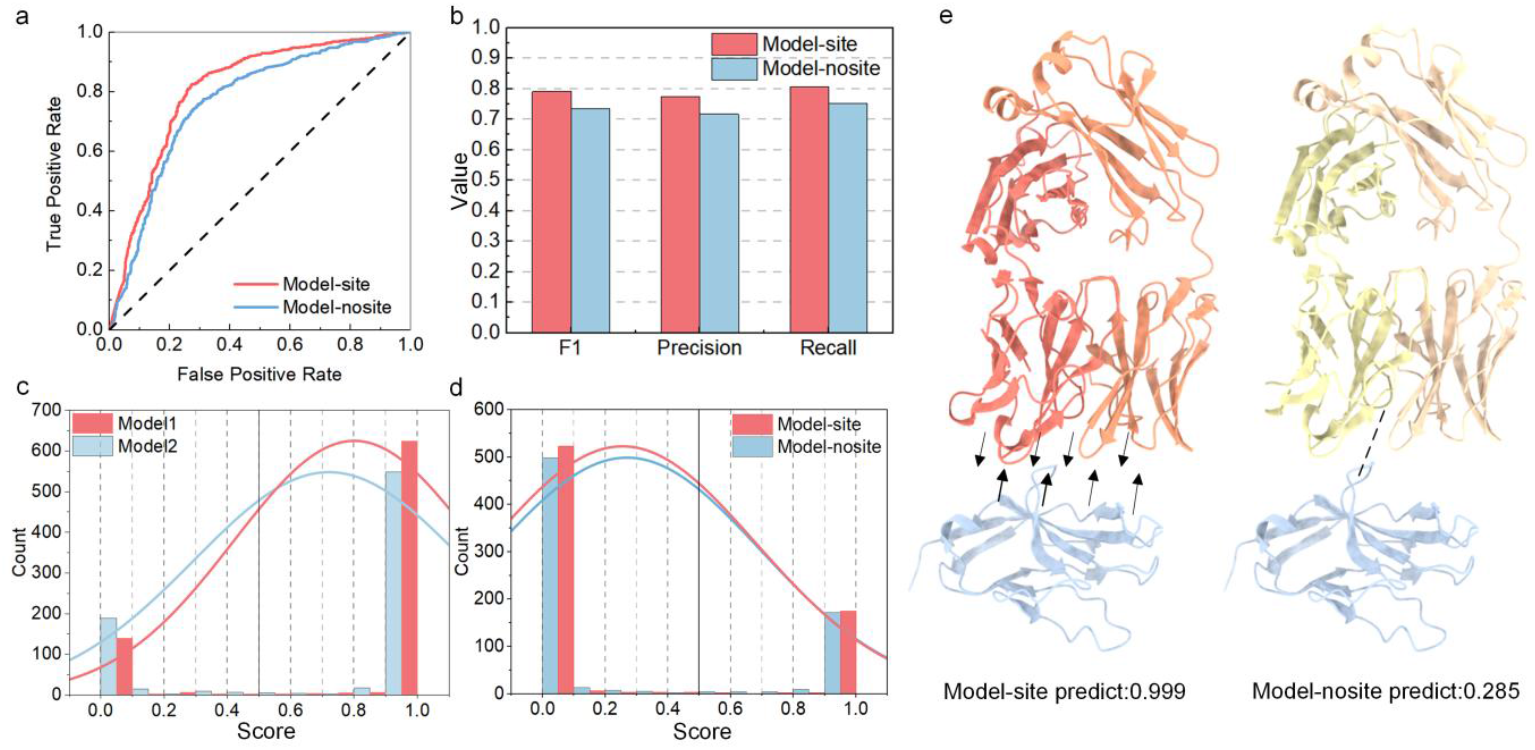
ProSiteHunter enhances antibody-antigen interaction prediction performance. **a**, Comparison of ROCAUC curves before and after the addition of predicted sites information. **b**, Describes the performance of the two models on F1, Precision and Recall. **c**, Distribution plot illustrates the classification performance of positive samples before and after incorporating binding site information. **d**, Distribution plot shows the classification performance of negative samples before and after the incorporation of binding site information. **e**, Schematic illustration of the complex (PDB ID: 6NMV) between signal regulatory protein α (SIRP α, blue) and Fab 218 antibody (orange/yellow) highlighting the difference between low and high predicted interaction probabilities.

To further validate the enhanced effect of integrating ProSiteHunter-predicted sites into MultisAAI^50^, we analyzed two representative case studies. First, Signal Regulatory Protein α (SIRPα ) is an inhibitory receptor expressed on the surface of myeloid cells, such as macrophages and neutrophils. By binding to the “don’t eat me” signal CD47, it inhibits the tumor-killing activity of myeloid cells. Kick-Off Fab115, an anti-SIRPα antibody, can effectively block this pathway^51^. As shown in **Fig. 6e**, incorporating site information increased the interaction probability between SIRPα and Kick-Off Fab115 from 0.285 to 0.999. Second, Hepatitis C Virus (HCV) infects approximately 1%-2% of the global population and is a leading cause of liver failure and hepatocellular carcinoma. The development of an effective HCV vaccine requires targeting conserved epitopes recognized by broadly neutralizing antibodies (AR3B)^52^. With ProSiteHunter’s precise epitope prediction, the probability of interaction between genotype 6a E2 protein and AR3B increased from 0.184 to 0.997 (**Supplementary Fig. 5**). Together, these cases illustrate that incorporating ProSiteHunter’s epitopes information can substantially improve antibody-antigen interaction modeling and screening, thereby providing valuable support for antibody design and related studies.

Overall, ProSiteHunter demonstrates excellent adaptability and transferability in antibody-antigen recognition tasks. The epitope information predicted by the model significantly improves the discriminative performance of downstream models, achieving comprehensive improvements in overall metrics while demonstrating higher sensitivity and reliability in distinguishing positive from negative samples. Further case studies confirm this, demonstrating that the key epitope regions captured by ProSiteHunter are highly consistent with the true binding interface and can accurately characterize the distribution characteristics of antigen binding sites. These results demonstrate that ProSiteHunter not only has powerful epitope mapping capabilities but also provides high-quality semantic representations to support the screening of antibody-antigen interactions.

## Discussion

Existing methods lack a general framework that can cover multiple types of binding sites. To address this challenge, we developed ProSiteHunter, a three-track semantic parsing network that includes forward-backward semantic analysis, local semantic extraction, and long-range semantic learning modules. By dynamically weighting and integrating multiple feature sources, the framework is able to more comprehensively characterize residue-residue interactions within a sequence. Experimental results show that ProSiteHunter can operate efficiently in various binding site prediction tasks and exhibits strong scalability. Although a few predicted residues may not correspond to the actual binding sites, its high throughput and low energy consumption make it an effective tool for the preliminary screening of functional binding sites.

ProSiteHunter further demonstrates its potential in modeling flexible regions and capturing dynamic conformational effects (**Figs. 3e and 5e**). This method can identify local flexible sites and binding site differences caused by dynamic conformational changes, providing a feasible strategy for analyzing flexible epitopes and dynamic structures. These results show that ProSiteHunter is highly complementary to structure-based methods, and the combination of the two is expected to provide a more comprehensive analytical perspective in complex systems, thereby improving the reliability and applicability of binding site predictions and helping to reveal protein-macromolecule interaction mechanisms that are difficult to elucidate with other methods.

As training data continues to grow and diversify, ProSiteHunter’s generalization capabilities are expected to further enhance. Not only will it maintain stable performance across a wider range of protein systems, but it will also naturally extend to the prediction of protein-small molecule ligand and protein-ion binding sites, further expanding its application. Furthermore, by combining ligand features with sequence information, ProSiteHunter is expected to achieve even higher accuracy and robustness in binding mode prediction. These improvements will provide a more reliable technical path for identifying functional binding sites, potentially making it a powerful tool for affinity prediction, molecular docking, and mutation effect analysis. This will facilitate a deeper understanding of protein binding mechanisms and hold great promise for applications in areas such as antibody design and antiviral vaccine development.

## Methods

### Data set

Protein-nucleic acid binding sites dataset: The DNA/RNA binding proteins were collected from the BioLiP database, released on December 5, 2018. If the minimum atomic distance between the target residue and the nucleic acid molecule is less than 0.5 Å plus the sum of the van der Waals radii of the two closest atoms, it is defined as a binding residue. Based on the release date, protein chains published before January 6, 2016, were assigned to the original training set (comprising 6731 DNA-binding protein chains and 6426 RNA-binding protein chains), while the remaining protein chains were allocated to the original test set (comprising 2843 DNA-binding protein chains and 1267 RNA-binding protein chains). The prediction of DNA/RNA binding residues faces a data imbalance issue, where the number of DNA/RNA binding residues is significantly smaller than that of non-binding residues; thus, data augmentation was applied to the original training set. Chains with sequence identities > 0.8 and TM scores > 0.5 were clustered, and the annotations of protein chains in the same cluster were transferred to the chain with the largest number of residues. After the migration and combined annotations, CD-HIT was used to remove redundant protein chains to reduce the sequence identity in the training set to less than 30%. Ultimately, 573 DNA-binding protein chains and 495 RNA-binding protein chains were obtained as the training set. Protein chains from the original DNA/RNA binding test set that shared over 30% sequence identity with any chains in the DNA/RNA binding training set were removed, resulting in 129 DNA-binding proteins and 117 RNA-binding proteins for the DNA and RNA binding test set. Please refer to GraphBind^17^ for the detailed processing procedure.

Protein-protein interaction sites dataset: We used the dataset used in EnsemPPIS^29^. This dataset is generated by combining three benchmark datasets, namely Dset_186^53^, Dset_72^53^, and PDBset_164^54^. The dataset comprises 422 non-redundant protein structures, each determined at a resolution better than 3.0 Å and sharing less than 25% sequence identity. After removing sequences longer than 2000 residues, 421 entries remained. If the absolute solvent accessibility of an amino acid decreases by at least 1 Å before and after protein binding, it is defined as an interaction site. The dataset is divided into a training set of 351 sequences (50 sequences are randomly selected from the training set as the validation set) and a test set (70 sequences).

Antibody-antigen conformational epitopes Dataset: The process of generating the conformational epitopes dataset for this dataset includes the following steps: The ANARCI^55^ tool is used to screen published protein structural sequences in the PDB database, including the heavy and light chains of Fab. Heavy/light Fab pairs are identified by calculating the distance between heavy and light chain subunit residues, and only those heavy and light chains that are in direct contact at a distance of 4.5 Å within a non-CDR region are considered heavy/light pairs. The CDR loop is defined using the Chothia number based on the notes in the ANARCI^55^ tool. A protein subunit that is not annotated as an antibody but has at least five residues interacting with antibody residues with L1/L2/L3 or H1/H2/H3 CDR antibody loops within a 4.5 Å radius is considered an antigen. Antigenic residues with a distance radius of less than 4.5 Å are defined as epitopes. According to the structure publication date, the dataset is divided into training set and test set. This test set includes structures first published in the PDB database after January 1, 2020, with no sequences of >70% available homologues prior to this date. The test set had a total of 101 antigen sequences, while the remaining antigens were divided into training set (713 items) and validation set (70 items). See SEMA 1.0^36^ for details.

### Sequence features

Protein language model: (1) SiteT5 is an enhanced protein language model fine-tuned based on ProtT5-XL-UniRef50 specifically for protein binding sites prediction tasks. ProtT5-XL-UniRef50^27^ is a large-scale protein language model with T5-3B architecture obtained from self-supervised pre-training of 45 million protein sequences on the UniRef50 dataset. In order to improve the performance of site prediction, we used UniRef30 for multiple sequence alignment (MSA) generation, which preserves a wider range of sequence diversity, helps to capture deeper evolutionary signals, and improves the recognition of conserved regions. During the MSA generation process, we employed the HHblits^56^ tool with stringent filtering parameters (E value threshold 0.000001, deredundancy 99%, coverage 25%) to ensure the quality of alignment and retain the top 50 high-quality MSA results for each sequence (see **Supplementary Table 8** for details). During the fine-tuning phase, in order to retain as much of the original information learned in the ProtT5 pre-trained model as possible, we fine-tuned only the last four layers of its 24-layer decoder. The LoRA (Low Rank Adaptation) strategy was adopted during fine-tuning to ensure efficient parameter updates while avoiding destabilizing the pre-trained representation. The final total number of fine-tuned parameters was 655,360. We searched for the corresponding MSAs using the protein sequences in their respective training sets for four different binding sites prediction tasks, and input these MSAs as important information into the model for 30 epochs to enable the model to learn specific patterns and characteristics of the protein family. To prevent overfitting, we used protein data with less than 30% similarity as a validation set, and ultimately obtained four high-performance fine-tuned models dedicated to different binding sites prediction tasks. Each of these amino acids is expressed using a tensor of 1024 dimensions. (2) ProstT5^41^ (Protein Structure-Sequence T5) uses the 3Di coding scheme proposed by Foldseek^42^ to convert the three-dimensional spatial structure information of proteins into one-dimensional discrete labeled sequences, so as to realize the bidirectional conversion between protein sequences and three-dimensional structures. The model is based on ProtT5-XL-UniRef50 and fine-tuned with 17M high-quality protein sequence-structure pairs (3D structure prediction from AlphaFoldDB^57^). Each of these amino acids is expressed using a tensor of 1024 dimensions. BLOSUM62^34^: It is based on evolutionarily conserved statistical properties and is used to measure the probability of substitution between different amino acids in protein sequence alignment. To avoid over-reliance on highly similar sequences, BLOSUM62 clusters sequences with a similarity of ≥62% (known as ‘62% clustering’). Conserved properties reflect the critical role of these regions in protein function or structure, so these conserved regions can be better described by BLOSUM62. It is of great significance for site identification.

Physicochemical properties: To provide a more specific description of amino acids, we introduce five physicochemical properties related to the site prediction task. Among these, the steric hindrance parameter measures the spatial volume or steric hindrance of the amino acid side chains, which affects spatial compatibility in protein interactions. The polarizability parameter describes the ease with which the electron cloud of the amino acid side chain deforms in an electric field, which is related to the solvent accessibility of proteins and influences intermolecular interactions. The volume parameter reflects the van der Waals volume occupied by amino acid side chains, providing insight into steric constraints in protein structures. The hydrophobicity of amino acids, reflecting their tendency to exclude water and aggregate in non-polar environments, can be leveraged to identify residues likely exposed on the protein surface. The isoelectric point parameter is used to describe the pH value at which the net charge of an amino acid or protein is zero, indirectly indicating the charge characteristics of the amino acid. A sequence of length L corresponds to a tensor representation of its physicochemical properties as (L,5).

Relative solvent accessibility characteristic is the ratio of the solvent accessible surface area of an amino acid residue in a protein structure to the maximum accessible surface area of that amino acid in a fully stretched ‘free’ state (e.g., Gly-X-Gly tripeptide conformation). This feature allows for a clear distinction between surface residues and buried residues, as the sites are present on the protein surface, and the addition of this feature narrows the scope of binding sites for prediction tasks. In addition, we believe that proteins are flexible and more likely to interact with other macromolecules. Therefore, secondary structure information is also necessary, so we have selected eight more detailed secondary structures to describe. These two characteristics were predicted by the NetSurfP3.0^58^ tool, which inputs the entire amino acid sequence to predict the probability of which secondary structure each amino acid belongs to in the sequence, as well as relative solvent accessibility information.

Amino acid propensity: Through a systematic statistical analysis of regions marked as binding sites in the training set, we found significant differences in amino acid composition between different types of protein binding sites. The binding sites of proteins to DNA/RNA exhibited highly similar amino acid distribution characteristics, with the positively charged hydrophilic amino acids lysine (K) and arginine (R) showing considerable enrichment, with their occurrence frequencies reaching 2.6 times and 3 times the average of twenty amino acids, respectively. Further analysis revealed that aspartic acid (D), glutamic acid (E), lysine (K), arginine (R), and leucine (L) were more abundant at protein-protein interaction sites, while antibody-antigen binding sites also displayed a high proportion of glycine (G), serine (S), and threonine (T). It is noteworthy that among these amino acids with a high proportion, leucine (L) is the only strong hydrophobic amino acid; the remaining are all hydrophilic, and more than half are charged amino acids (D/E/K/R), suggesting that electrostatic interactions and hydrophilicity may play an important role in sites recognition. The full statistical results can be found in **Supplementary Fig. 2**.

After that, we have added a symmetry normalized position code that measures the degree of symmetry of amino acids in the sequence, with outputs ranging from [0, 1]. It can be used in protein sequence analysis to enhance a model’s understanding of the global structure.

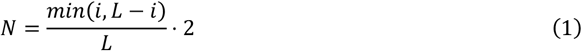

*i* represents the index of residues, while *L* denotes the total length of the protein sequence.

### Network architecture

We adopt an encoder-decoder architecture, which primarily comprises two components: a Multi-Source Feature Fusion (MSFF) module and a Multi-Level Interaction Learning (MIL) module. Below, we provide a detailed description of these modules. While various types of features are introduced to characterize protein sequences, a key challenge lies in effectively processing these heterogeneous features. To address this, we designed the MSFF module, which consists of three parts: a Multi-Scale CNN module, a BiLSTM module, and a Gated Attention module. The tensors representing different patterns learned by these three networks are then mapped as Queries (Q), Keys (K), and Values (V) and fed into a cross-attention layer for integrative information fusion. In order to simplify the representation of features, we refer to all other features, except for the language model features SiteT5 and ProstT5, as the protein composite features (pcf).

Let us first introduce the multi-scale convolution module. To better learn the relationships between local residues, we designed the multi-scale convolution module. Conv1D mainly captures the contextual representations of residues with local biases and learns global protein features by assembling the local features of all residues. By aggregating the residue representations through convolutional kernels of different sizes, specifically 3, 5, and 7, we can learn information across different receptive fields.

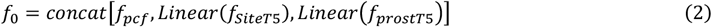

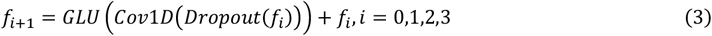

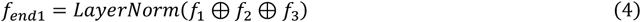

Among these, *f*_*pcf*_ represents a 43-dimensional feature, *f*_*SiteT*5_ is a 1024-dimensional feature that is transformed into a 102-dimensional feature after passing through a linear layer, and *f*_*prostT*5_ is a 1024-dimensional feature that is reduced to a 102-dimensional feature after a linear layer. *f*_0_ contains features obtained from multi-scale convolution, while *f*_*i*_, *i* = 0,1,2,3 represents feature tensors after applying different convolution kernels (3, 5, 7). ⊕ describes the serial relationship, and *f*_*end*1_ is a 64-dimensional tensor.

The BiLSTM module simultaneously includes a forward LSTM (which processes sequences from past to future) and a backward LSTM (which processes sequences from future to past), ultimately concatenating the hidden states from both directions. This module is capable of capturing past (historical) information while also leveraging future (subsequent) information, selectively memorizing important data (such as features at critical time steps) and ignoring irrelevant data. It is particularly well-suited for handling long sequence data, which is why we have incorporated this module.

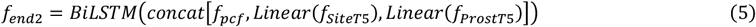

To effectively capture the interaction information between long-distance residue pairs in protein sequences and highlight the important residue tensors that contribute significantly to structure or function, we have employed a gated attention module. The attention module is proficient at capturing long-range information; however, it is unable to dynamically adjust the focus on important features. Through gated units, high-weight residue signals are preserved and enhanced via gating (such as potential residue sites).

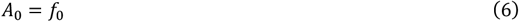

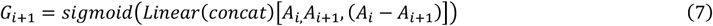

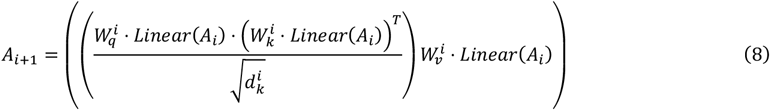

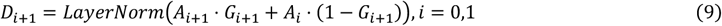

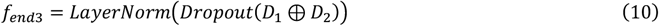

Where *A*_0_ is the feature of the input, which is clearly described in multiscale convolution. *A*_1_, *A*_2_ is the tensor after passing through the 8-head self-attention network, *G*_1_, *G*_2_ is the weight generated by the gated network is a 1-dimensional tensor, *D*_1_, *D*_2_ is the tensor output after the gated weights are multiplied by points, ⊕ describes the concatenation relationship, and *f*_*end*3_ is a 64-dimensional tensor.

Three networks are then mapped as Queries (Q), Keys (K), and Values (V) and fed into a cross-attention layer for integrative information fusion.

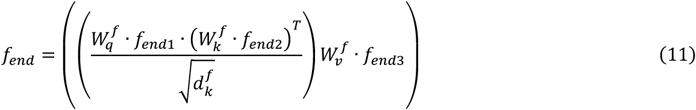

*f*_*end*_ is the tensor after the 8-head cross-attention module.

The decoder is designed as a Multi-Layer Interaction Learning (MIL) module composed of three identical stacked sub-blocks. Each sub-block consists of a gated multi-head self-attention unit and a position-wise feed-forward network (PFFN), enabling the MIL to iteratively aggregate and enhance critical feature information. The integrated features are subsequently passed through a multilayer perceptron (MLP) to produce the final prediction output.

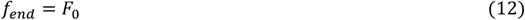

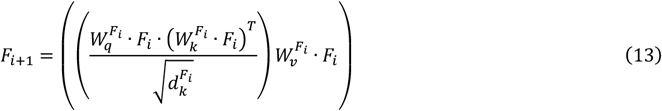

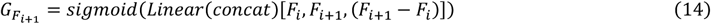

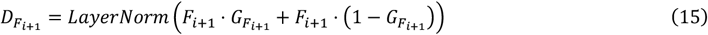

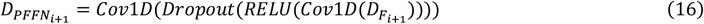

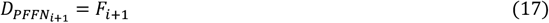

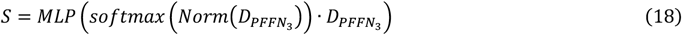

*F*_0_ is the tensor after the 8-head cross-attention module, *F*_*i*_, *i* = 0,1,2 is the tensor after the 8-head self-attention, 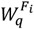 represents the weight matrix, 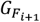 is the gating parameter, and 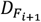 is the tensor output after the gating layer. 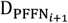 is the tensor output from position-wise feed-forward network and *S* is the result of the MLP output.

### Network design intent

Multi-Source Feature Fusion module (MSFF): In this tri-track semantic parsing structure, the Query (Q), Key (K), and Value (V) components serve distinct informational roles: Q determines what the model should focus on, K provides the contextual semantics to be referenced, and V defines the information content to be aggregated and updated in the final representation. Therefore, their corresponding subnetworks are designed to be semantically complementary and functionally aligned. Specifically, the Multi-Scale CNN captures local sequence patterns and short-range residue dependencies with different receptive fields, providing high-resolution and position-sensitive features. Hence, it is most suitable as Q (Query), initiating attention queries toward relevant contexts. The BiLSTM module bidirectional contextual dependencies, offering a smooth and consistent global semantic reference, thus serving as K (Key) to provide globally aligned semantic coordinates for each query. Meanwhile, the Gated Attention module performs global aggregation and adaptive weighting, suppressing irrelevant noise while amplifying functionally important signals; it is therefore assigned as V (Value), providing the information that will be integrated during fusion. After mapping to Q, K, and V spaces, these representations are fed into a cross-attention module, where attention weights are computed to dynamically align heterogeneous feature spaces and achieve context-sensitive fusion. This hierarchical information flow (from local feature extraction (Q) to global semantic alignment (K) and final information aggregation (V)) ensures explicit semantic propagation and functional complementarity within the model. Reversing this correspondence would disrupt the intrinsic logic of the attention mechanism, weakening both the interpretability and the effectiveness of the feature fusion process. Multi-Level Interactive Learning (MIL): Although static feature fusion can integrate multi-modal semantics, it remains insufficient for capturing the dynamic coupling and cooperative effects among residues. To address this limitation, the MIL module adopts a hierarchical three-block architecture, where each block consists of an eight-head self-attention mechanism, a gating mechanism, and a position-wise feed-forward network (PFFN).Within each block, the multi-head self-attention mechanism models residue relationships across multiple semantic subspaces in parallel, thereby implicitly capturing multi-scale dependencies ranging from local to long-range interactions and enabling dynamic context aggregation. The gating mechanism adaptively modulates attention weights according to feature relevance, suppressing noisy activations while emphasizing functionally critical residues. The subsequent PFFN performs nonlinear transformation and semantic enhancement on the aggregated features to improve discriminative capacity and generalization. Through progressive three-layer interaction, the MIL module achieves hierarchical semantic evolution (from local dependency learning to global cooperative modeling) enabling ProSiteHunter to capture high-order residue interactions directly from sequence data and deliver superior accuracy, robustness, and biological interpretability in binding-site prediction.

### Network hyperparameters

All network parameters are as follows: In the multi-source feature fusion module, sequential features are processed by three distinct networks, where we have set up two LSTM layers, each containing 32 units. To prevent overfitting, a dropout probability of 0.3 is applied between the layers. As this is a bidirectional LSTM, the output concatenates the hidden states from both the forward and backward directions, resulting in a tensor of dimension (L, 64). For the multi-scale convolution module, we have configured three layers of 1D convolutional layers with kernel sizes of 3, 5, and 7. Each convolutional layer leverages residual connections to enhance network stability, producing a tensor of dimension (L, 64). The gated attention module comprises two concatenated layers of eight-head self-attention, with an inter-layer dropout probability of 0.1. Additionally, gated parameters generated through softmax are set to dynamically adjust in between layers, resulting in a tensor of dimension (L, 64). The features outputted by the three networks are multiplied by their corresponding weights to be reused as q, k, and v for attention learning. Finally, the MLP includes three hidden layers of fully connected networks.

### Training

ProSiteHunter is implemented in Python in Pytorch and trained for up to 30 epochs. Since it is a common sequence-based framework, it consumes less resources and the training batch size is 1. The Radam optimizer is used to minimize the loss, and the loss function chosen is a weighted cross-entropy loss function designed to address unbalanced sample data, where class 0 has a weight of 1 and class 1 has a weight of 5.

Given the input samples *X* ∈ *R*^*L*×*d*^, (L is the number of residues and d is the feature dimension), the model outputs the raw score (logits) *Z* ∈ *R*^*L*×2^ through MLP, where *z*_*i*,0_ represents the score of the ith and the sample belongs to category 0, and *z*_*i*,1_ represents the score that the ith sample belongs to the positive class 1. Convert logits to probability distributions via the softmax function.

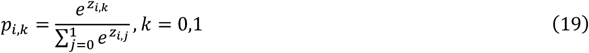

where *p*_*i*,0_ and *p*_*i*,1_ represent the predicted probability that residue i belongs to the negative class and the positive class respectively, and satisfy *p*_*i*,0_ + *p*_*i*,1_ = 1.

In order to deal with the category imbalance problem, a higher weight *w*_1_ = 5 is given to the positive class (category 1), and the weight *w*_0_ = 1 is given to the negative class (category 0). The loss function is defined as:

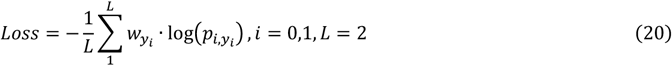

where *y*_*i*_ = 0,1 is the true label of residue i, and the optimization goal is to minimize Loss.

The learning rate is initially set at 0.0005 and gradually decreases over the course of training, following an exponential decay policy outlined. Using an NVIDIA GeForce RTX 3090 GPU to train, it takes about 1 hours to train the network.

## Reporting summary

Further information on research design is available in the Nature Portfolio Reporting Summary linked to this article.

## Data availability

The authors declare that the data supporting the results and conclusions of this study are available within the paper and its Supplementary Information.

## Code availability

ProSiteHunter is available for download via GitHub at https://github.com/iobio-zjut/ProSiteHunter/tree/main.

## Acknowledgements

We thank members of the Guijun Zhang lab for discussion and feedback. Computational resources were provided by the College of Information Engineering at Zhejiang University of Technology. This work was supported by the National Key R&D Program of China [2022ZD0115103, G.Z.], the National Nature Science Foundation of China [62173304, G.Z.; 62203389, X.Z.; 62573386, G.Z.], the “Pioneer” and “Leading Goose” R&D Program of Zhejiang [2025C01190, G.Z.], the Zhejiang Province High-level Talent Special Support Program [2023R5248, G.Z.], and Fundamental Research Funds for the Provincial Universities of Zhejiang [RF-C2024006, X.Z.].

## Competing interests

The authors declare no competing interests.

## Additional information

**Correspondence** and requests for materials should be addressed to G.Z.

## Supplementary Information

### Supplementary Tables

**Supplementary Table 1.** Detailed description of input features for the network.

| Feature |  | Description | Dimensions |
| --- | --- | --- | --- |
| Evolutionary features | SiteT5 | SiteT5 is an enhanced protein language model that retains the first 50 MSA data per sequence for fine-tuning of ProtT5-XL-UniRef50 | L×1024 |
|  | ProstT5 | ProstT5 selected 17M high-quality protein sequence-structure pairs and fine-tuned ProtT5-XL-UniRef50 via a 3Di coding format | L×1024 |
| Statistical features | BLOSUM62 Matrix | Blosum62 is based on evolutionarily conserved statistical properties and is used to measure the probability of substitution between different amino acids in protein sequence alignment. | L×24 |
|  | Amino acid propensity | Through systematic statistical analysis of the regions labeled as binding sites in the training set, we found significant differences in amino acid composition between different types of protein binding sites. | L×1 |
|  | Physicochemical properties | Five physicochemical properties relevant to site prediction tasks. steric hindrance parameters, polarizability, amino acid side chain volume, hydrophobicity, and isoelectric point | L×5 |
| Geometric features | secondary structure | The flexible loop regions are more likely to interact with other macromolecules. Therefore, eight more detailed secondary structures have been selected for description. | L×8 |
|  | Positional encoding | A symmetrical normalized position code that measures the degree of symmetry of amino acids in a sequence. | L×1 |
|  | Relative solvent accessibility | Relative solvent accessibility can clearly distinguish between surface and buried residues, and the addition of this feature narrows the range of binding sites for prediction tasks. | L×1 |

**Supplementary Table 2.** The number of positive and negative samples in the protein-DNA binding site dataset and the protein-RNA binding site dataset.

| **Datasets** | **protein-DNA**<br>**(train)** | **protein-DNA**<br>**(test)** | **protein-RNA**<br>**(train)** | **protein-RNA**<br>**(test)** |
| --- | --- | --- | --- | --- |
| binding sites | 14479 | 2240 | 14609 | 2031 |
| non-binding sites | 145404 | 35275 | 122290 | 35314 |

**Supplementary Table 3.** Detailed description of characteristic ablation experiments.

|  |  |
| --- | --- |
| **Module1#** | ProstT5 |
| **Module2#** | ProstT5+SiteT5 (Evolutionary features) |
| **Module3#** | Evolutionary features+Statistical features |
| **Module4#** | Evolutionary features+Statistical features+Physicochemical |
| **Module5#** | Evolutionary features+Statistical features+ Physicochemical+Geometric features |

**Supplementary Table 4.** Compare the performance of ProSiteHunter, CLAPE-DB, iDRNA-ITF, and DRNApred on the protein-DNA binding site test set (Test129).

| **Method** | **ROCAUC** | **PRAUC** | **F1** | **MCC** |
| --- | --- | --- | --- | --- |
| ProSiteHunter | 0.911 | 0.496 | 0.485 | 0.473 |
| CLAPE-DB | 0.865 | 0.394 | 0.380 | 0.367 |
| iDRNA-ITF | 0.834 | 0.400 | 0.421 | 0.386 |
| DRNApred | 0.692 | 0.143 | 0.214 | 0.150 |

**Supplementary Table 5.** Compare the performance of ProSiteHunter, CLAPE-RB, iDRNA-ITF, and DRNApred on the protein–RNA binding site test set (Test117).

| **Method** | **ROCAUC** | **PRAUC** | **F1** | **MCC** |
| --- | --- | --- | --- | --- |
| ProSiteHunter | 0.852 | 0.302 | 0.327 | 0.316 |
| CLAPE-RB | 0.788 | 0.179 | 0.281 | 0.240 |
| iDRNA-ITF | 0.702 | 0.186 | 0.276 | 0.231 |
| DRNApred | 0.662 | 0.124 | 0.175 | 0.122 |

**Supplementary Table 6.**
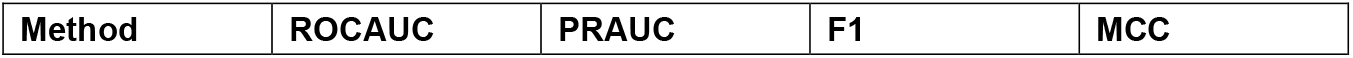
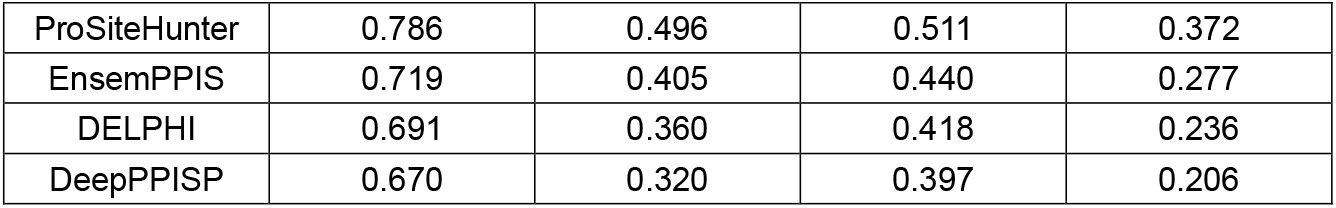
Compare the performance of ProSiteHunter, EnsemPPIS, DELPHI, and DeepPPISP on the protein-protein binding site test set (Test70).

**Supplementary Table 7.** Compare the performance of ProSiteHunter, CALIBER, Bepipred-3.0, and SEMA2.0-1D on the antibody-antigen binding site test set (Test101).

| Method | ROCAUC | PRAUC | F1 | MCC |
| --- | --- | --- | --- | --- |
| ProSiteHunter | 0.770 | 0.263 | 0.300 | 0.237 |
| CALIBER | 0.756 | 0.245 | 0.277 | 0.216 |
| Bepipred-3.0 | 0.733 | 0.211 | 0.272 | 0.214 |
| SEMA2.0-1D | 0.730 | 0.187 | 0.223 | 0.179 |

**Supplementary Table 8.** Analyze the impact of fine-tuning with sub-MSAs of varying depths on the performance of the protein–protein binding site prediction task.

| MSA | Sequence Number | ROCAUC | PRAUC | F1 | MCC |
| --- | --- | --- | --- | --- | --- |
| No fine-tuning | 0 | 0.764 | 0.454 | 0.487 | 0.338 |
| Top1 | 351 | 0.775 | 0.488 | 0.499 | 0.355 |
| Top25 | 7436 | 0.780 | 0.492 | 0.503 | 0.360 |
| Top50 | 14690 | 0.786 | 0.496 | 0.511 | 0.372 |
| Top75 | 21850 | 0.785 | 0.496 | 0.507 | 0.360 |
| Top100 | 28934 | 0.782 | 0.486 | 0.505 | 0.363 |
| Top200 | 56740 | 0.774 | 0.488 | 0.496 | 0.354 |

### Evaluation metrics

The protein-macromolecule binding sites prediction task is a binary classification task, therefore, we followed previous studies and used recall, precision, F1 score, and Matthew correlation coefficient (MCC) as metrics to evaluate the performance of our method. The calculation formula is as follows:

Recall: The proportion of positive examples correctly identified by the model to all positive examples.

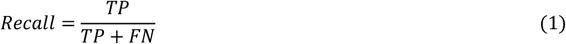

Precision: The proportion of positive cases predicted by the model to the true positive examples.

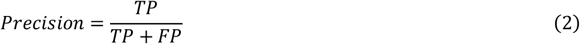

F1: The harmonized average of precision and recall.

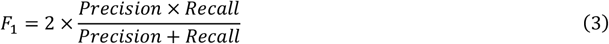

MCC: The correlation coefficient of all prediction results is combined, and the value range is [-1, 1], where 1 indicates a perfect prediction.

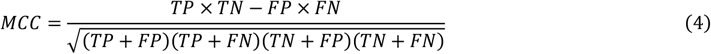

where TP, FN, TN, FP represent the number of true positives, false negatives, true negatives, and false positives, respectively. In particular, when the ratio of positive and negative samples is not well balanced, F1 and MCC are more objective indicators of model performance than recall or precision alone, as the binding sites are usually much smaller than the non-binding sites (positive and negative sample imbalance). In addition, we also used ROCAUC and PRAUC to evaluate the model from the perspective of global threshold changes, where PRAUC is more sensitive to class imbalance and is a key indicator of conjugation sites prediction.

## Supplementary Figures

**Supplementary Fig. 1.**
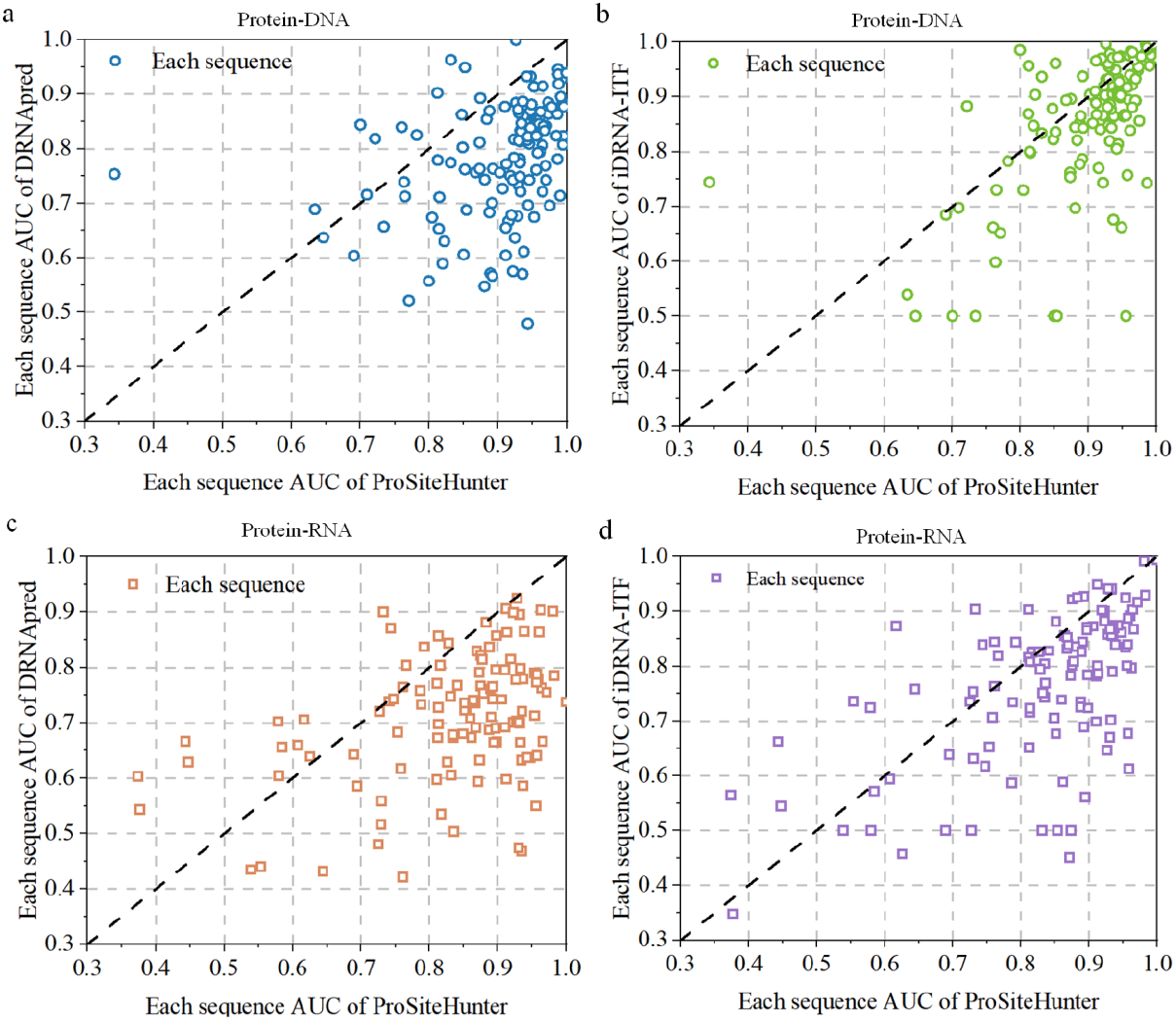
**a**, Scatter plots display the sequence-wise AUC values of ProSiteHunter versus DRNApred for protein-DNA binding. **b**, Scatter plots display the sequence-wise AUC values of ProSiteHunter versus iDRNA-ITF for protein-DNA binding. **c**, Scatter plots display the sequence-wise AUC values of ProSiteHunter versus DRNApred for protein-RNA binding. **d**, Scatter plots display the sequence-wise AUC values of ProSiteHunter versus iDRNA-ITF for protein-RNA binding.

**Supplementary Fig. 2.**
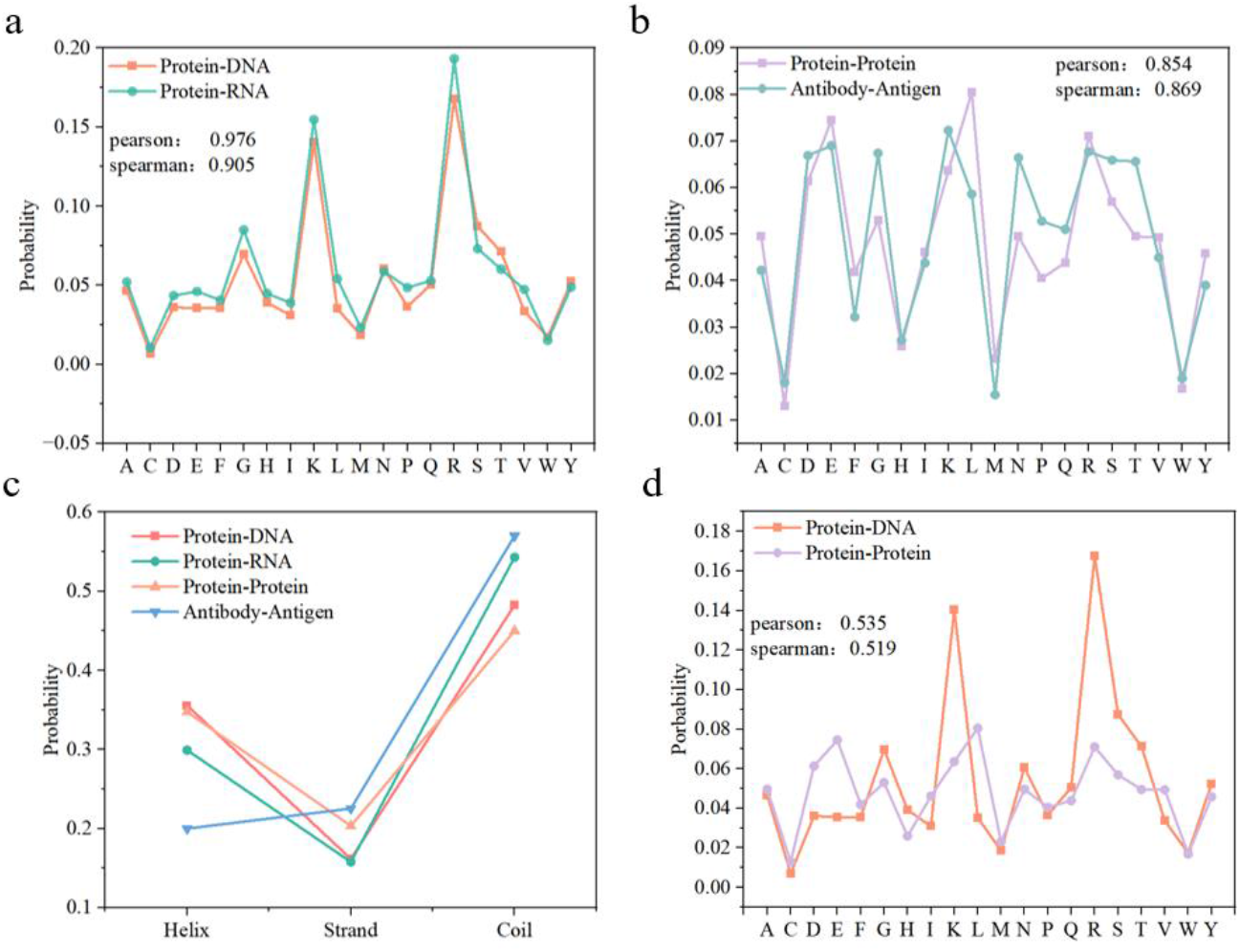
**a**, Amino acid frequency statistics at protein-DNA binding sites and protein-RNA binding sites. **b**, Amino acid frequency statistics at protein-protein binding sites and antibody antigen binding sites. **c**, the secondary structure composition of protein-DNA, protein-RNA, protein-protein, and antibody-antigen binding sites. **d**, Amino acid frequency statistics at protein-DNA binding sites and protein-protein binding sites.

We then analyzed the amino acid trends for different types of binding sites statistically based on the training set data. The binding sites of DNA-binding proteins and RNA-binding proteins are highly consistent in amino acid composition, with a Pearson correlation coefficient of 0.976 and a Spearman correlation coefficient of 0.905, which may indicate that there is a similarity in the binding patterns between the two types of proteins and nucleic acid molecules. Similarly, the amino acid composition at the protein-protein interaction sites and the antibody antigen binding sites (epitopes) was highly similar (Pearson = 0.854, Spearman = 0.869), suggesting that despite the specificity of antibody antigens, their binding patterns are the same as those of conventional protein-protein interactions. Specific details will be provided in **Supplementary Figs. 1a, 1b**

In the task of predicting four types of binding sites (statistically based on the training set data), we found that the prediction difficulty of antibody-antigen epitopes and RNA-binding protein sites was significantly higher than that of DNA-binding protein sites and protein-protein interaction sites. This phenomenon seems paradoxical, as the previous analysis has shown that these binding sites are highly similar in amino acid composition (RNA-binding protein sites and DNA-binding protein sites, protein-protein interaction sites, and antibody-antigen epitopes) and should theoretically exhibit similar binding patterns. To explore the potential reasons for this difference, we conducted a more in-depth analysis of the data. **Supplementary Fig. 1c** statistically analyzed the secondary structure composition of protein-DNA, protein-RNA, protein-protein, and antibody-antigen binding sites, and the results showed that the loop was dominant among the four binding sites (accounting for 45%-57%), indicating that it plays a key role in the process of sites recognition. It is worth noting that the protein sites region that binds to RNA and the epitopes region that binds to antibody have the highest proportion of loops (54.3% and 57.2%, respectively), and these two types of binding sites also show higher prediction difficulty, which may indicate that the high flexibility of loops may be an important factor leading to the decrease in prediction accuracy. In addition, unlike the other three types of binding sites tasks, antigen sites (epitopes) exhibit a unique secondary structure distribution characteristic: the proportion of *β* fold is higher than that of *α* helix. We speculate that this phenomenon may be related to the fact that the hypervariable loop region of the epitopes accounts for nearly 60% of the total, and it needs to provide a stable structural framework *β* folding to maintain its conformational stability.

A comparative analysis of the amino acid tendencies of protein-DNA and protein-protein binding sites (Supplementary Fig. 1d) shows that the two classes of binding sites exhibit significantly different binding preferences (Pearson:0.515, Spearman:0.519), suggesting that different types of protein-macromolecule interactions may follow different recognition mechanisms.

**Supplementary Fig. 3.**
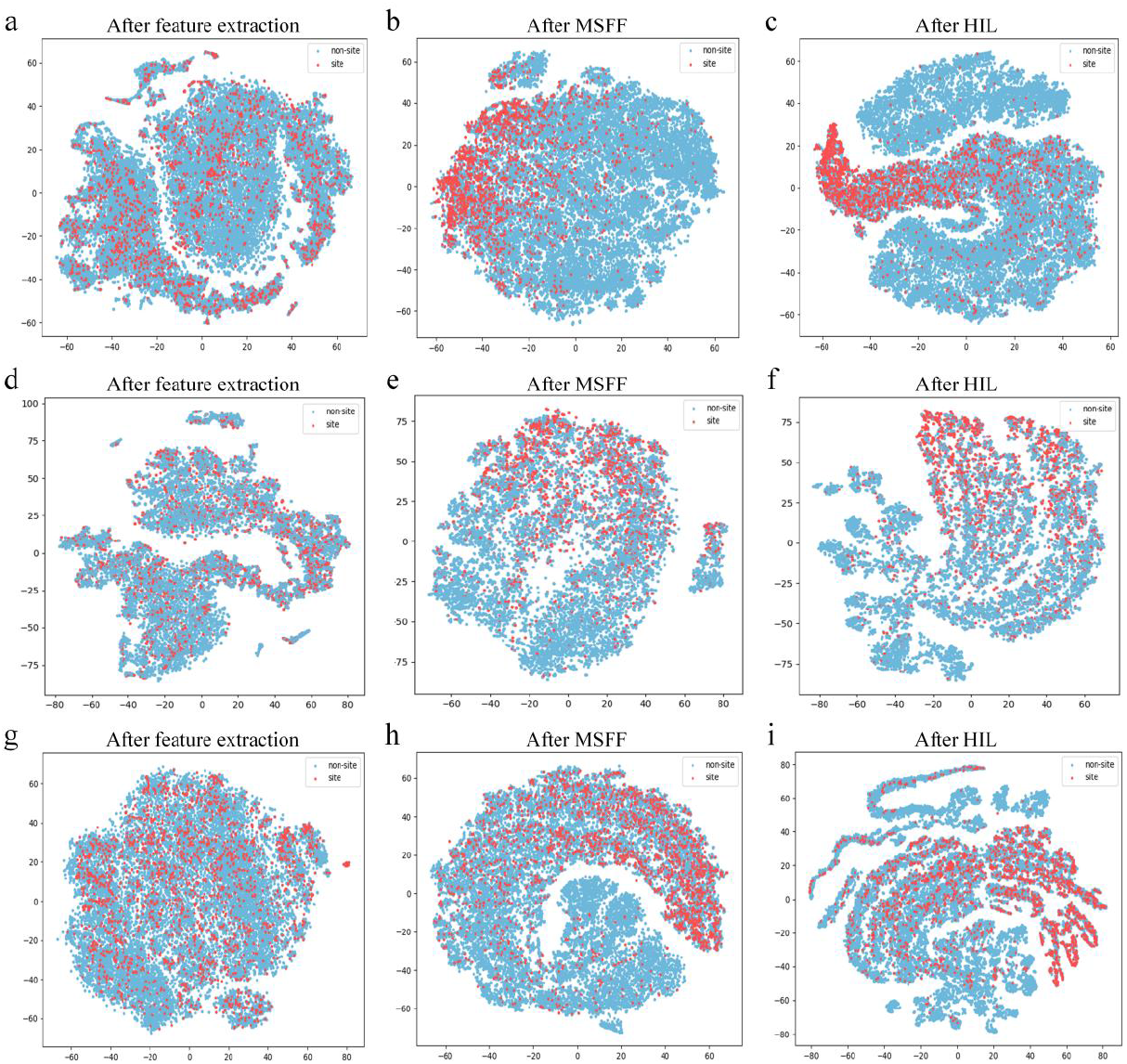
**a-c**, Use the t-SNE graph to map the high-dimensional tensor after feature extraction, MSFF module, and HIL module into a two-dimensional space to describe the relationship between protein-RNA sites and non-sites. **d-f**, Use the t-SNE graph to map the high-dimensional tensor after feature extraction, MSFF module, and HIL module into a two-dimensional space to describe the relationship between protein-protein sites and non-sites. **g-i**, Use the t-SNE graph to map the high-dimensional tensor after feature extraction, MSFF module, and HIL module into a two-dimensional space to describe the relationship between antibody-antigen sites and non-sites.

**Supplementary Fig. 4.**
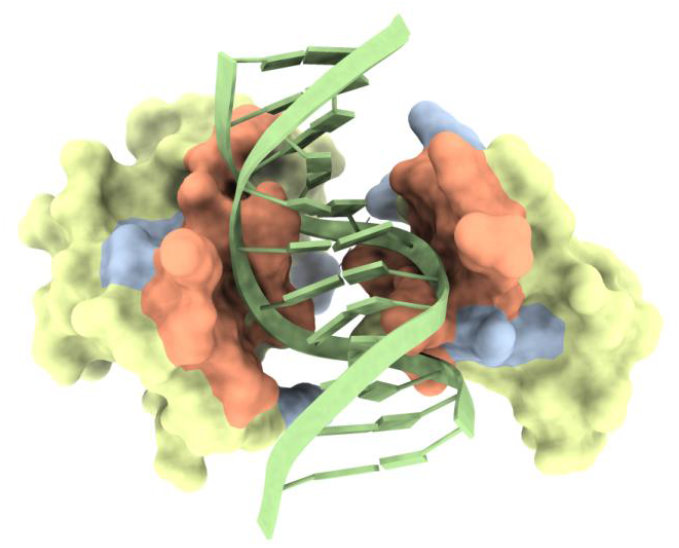
This protein (PDB ID: 5FD3) is a complex of DNA (green) and the Lin54 tesmin domain (yellow). The orange portion represents the actual binding site, while the blue and orange regions represent the sites predicted by ProSiteHunter.

**Supplementary Fig. 5.**
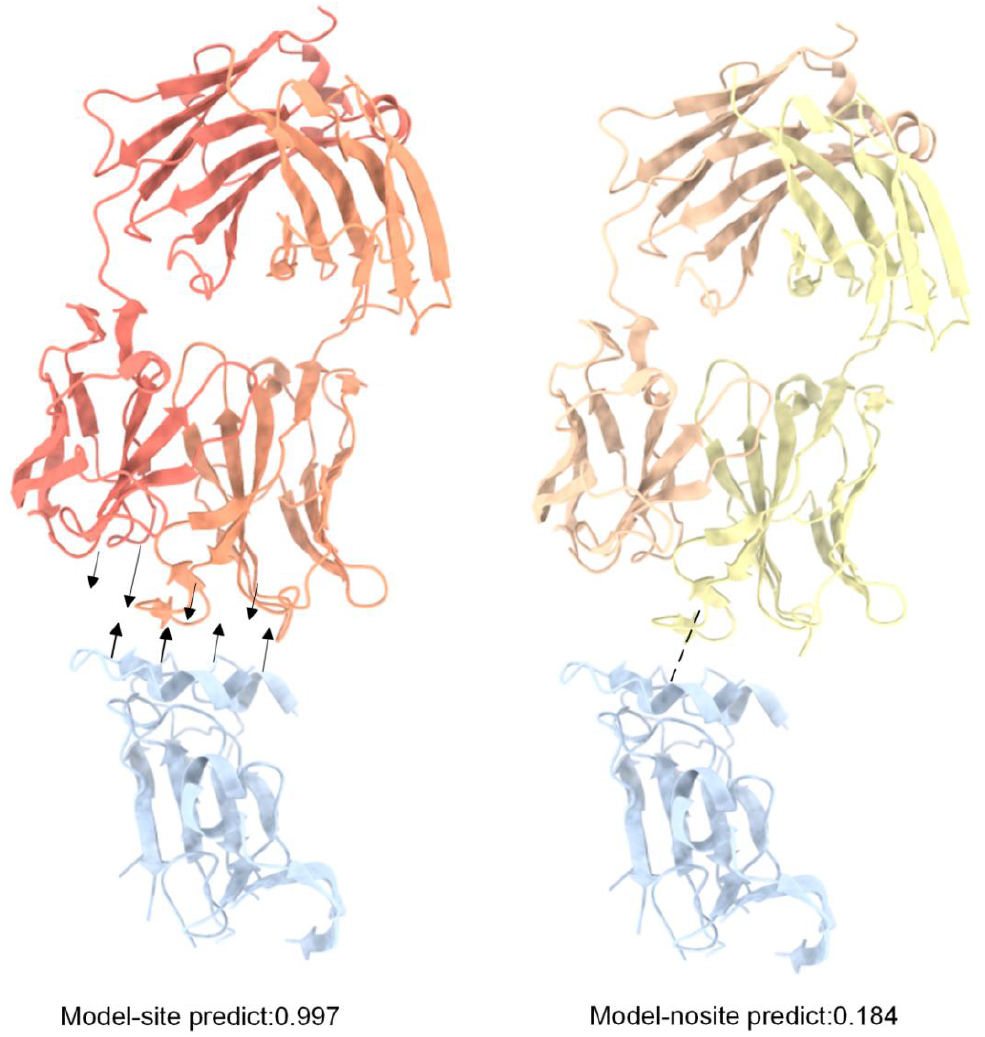
Schematic diagram of the structure(PDB ID: 6BKC) of the HCV envelope glycoprotein E2 core (from genotype 6a, blue) in complex with the broadly neutralizing antibody AR3B (orange/yellow) highlighting the difference between low and high predicted interaction probabilities.

## Notes

### Competing Interest Statement

The authors have declared no competing interest.

